# Self-amplifying RNA enables rapid, durable, integration-free programming of hiPSCs

**DOI:** 10.1101/2025.10.24.684179

**Authors:** Catherine M. Della Santina, Deon S. Ploessl, Nicole Lindsay-Mosher, Caroline G. Brown, Mary E. Ehmann, Albert Blanch-Asensio, Eung Chang Kim, Edward S. Boyden, Kate E. Galloway, Laurie A. Boyer

## Abstract

Genetic modification of human induced pluripotent stem cells (hiPSCs) is a powerful approach to measure and manipulate the cellular processes underlying differentiation and disease. Conventional genetic engineering of hiPSC lines requires a laborious process involving transfection, selection and expansion that can result in karyotypic abnormalities or transgene silencing during differentiation, limiting their applications. Self-amplifying RNA (saRNA) delivery is a potential alternative integration-free method for durable expression of transgenes. Here, we used saRNA to deliver transcription factors and functional reporters in hiPSCs and demonstrate that expression can persist for weeks. Specifically, saRNA delivery enables highly efficient forward programming to Ngn2-induced neurons and enables measurement of functional reporters over time. We show that a single transfection of saRNA encoded jRCaMP1b reporter in hiPSCs generates sustained expression throughout differentiation to 3D cardiac spheroids. The persistence of the reporter allows measurement of calcium dynamics at a single-cell and population level over weeks, allowing tracking of cardiomyocyte maturation and drug responses. Together, our systematic analysis shows that saRNA provides sustained transgene expression in hiPSCs, supporting integration-free cell-fate programming and measurement of functional reporters in clinically relevant model systems.

**Highlights:**

- A single saRNA transfection generates durable transgene expression
- saRNA transfection of Ngn2 in hiPSCs results in robust neuronal differentiation
- saRNA-delivery of functional reporters enables single-cell analysis of primary and hiPSC-derived cells
- saRNA-based sensor allows monitoring of maturation and drug responses in 3D cardiac spheroids

## Introduction

Genetically encoded reporters and transcription factors are powerful tools for measuring and modifying cellular function and fate. Choice of transgene delivery method sets scalability, durability, and precision in cellular engineering. For example, transient delivery of RNA and plasmids supports rapid, integration-free expression, but it is not well suited for applications that require expression for multiple days or weeks. Alternatively, integration of transgenes using programmable nucleases, transposons, or lentiviruses can support durable expression but require time-consuming cell line engineering and selection strategies that may affect cellular physiology [1]. Moreover, extended culture as needed for isolation of clonal lines intensifies single-cell bottlenecks that can induce genetic drift, posing a challenge for the generation and comparison of multiple hiPSC lines [2]. Although landing pad hiPSC lines can streamline cell line construction, their initial generation and validation requires months, limiting the inherent scalability of this approach [3]. Moreover, integration approaches often result in transgene silencing, limiting the use of hiPSCs for long-term gene expression, therapeutic gene delivery, and other synthetic biology applications [5],[6]. While adenoviral-associated virus (AAV) addresses some of these challenges, the small packaging capacity of AAV prevents delivery of large cargoes [4] and production of AAV demands optimization which can limit scale-up [5]. Modified mRNA (modRNA) supports applications where transient expression is sufficient [1], [3], [4], [6], [7], [8], [9], but requires multiple transfections to achieve durable responses, limiting scalability and compromising cell viability [10], [11], [12], [13]. In contrast, self-amplifying RNA (saRNA) offers key advantages for delivering transgenes including higher expression at lower doses and durable expression [14] in applications including protein replacement therapy [15], hiPSC reprogramming [16], and inducible gene circuits [14], yet there has been limited systematic characterization of saRNA as a tool to manipulate hiPSC systems [14].

Delivery of transgenes using saRNA has been used in applications including protein replacement therapy [15], hiPSC reprogramming [16], and inducible gene circuits [17]. saRNAs are derived from positive-strand RNA viruses, encoding the viral conserved sequence elements (CSEs) and non-structural protein genes (nsP1-4) for replication. A transgene can be inserted as a single transcript replicon that facilitates self-replication within the cytoplasm of a host cell (**Figure 1A**). The endogenous subgenomic promoter (SGP) located upstream of the transgene is separately transcribed by nsP1-4, amplifying transgene transcript levels. Translational applications of saRNAs primarily focus on vaccine development for infectious disease and cancer [18]. In saRNA-based vaccines, the amplification of transgene expression allows saRNAs to produce similar levels of antigen compared to mRNA with a fraction of the dose [19]. Since saRNAs are replicated in the cytoplasm and do not contain double stranded DNA, there is a low risk of genomic integration [20]. Thus, saRNAs could be used more broadly as a powerful tool for durable, scalable, integration-free delivery of transgenes to hiPSCs [21].

**Figure 1.**
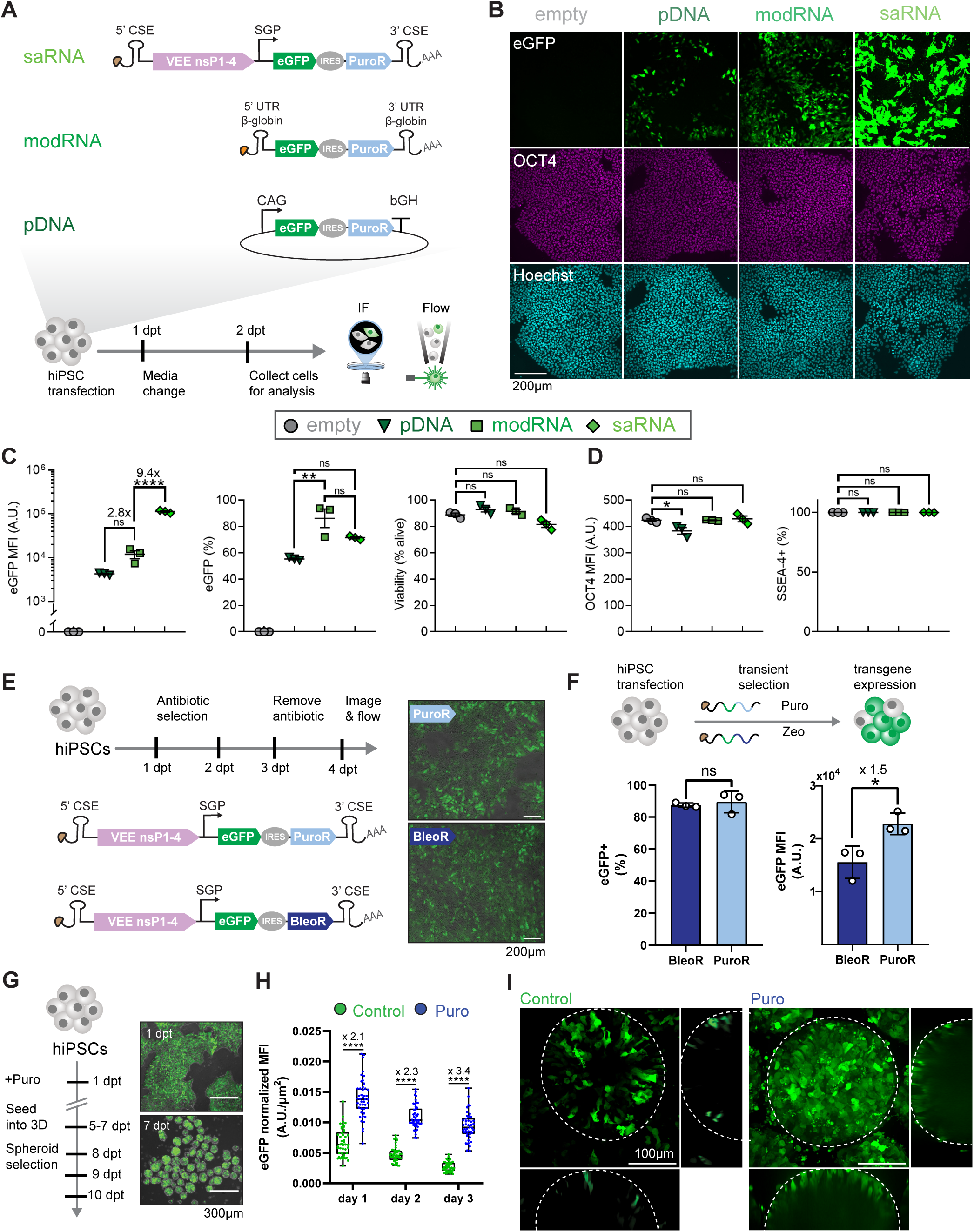
saRNA enables sustained transgene expression in 2D and 3D hiPSC cultures. Expression from saRNA confers selection marker resistance without impacting cell viability or pluripotency markers in hiPSCs. **A**. hiPSCs are transfected with equal amounts (by mass) of either saRNA, modRNA, or pDNA eGFP-IRES-PuroR transgenic cassette. For saRNA, co-opting the alphaviral regulatory sequences drives high expression of transgenes located downstream of the subgenomic promoter (SGP). For modRNA, B-globin UTRs enable effective translation of coding mRNA sequences. For pDNA, the CAG promoter and bovine growth hormone (bGH) polyadenylation signal allow efficient expression of transgenes in hiPSCs. **B**. Representative fluorescence microscopy images of hiPSCs fixed at 2 days post transfection (dpt) with each delivery construct in A. **C**. Flow cytometry analysis of eGFP expression (MFI: Mean Fluorescence Intensity, measured at 2 dpt in Arbitrary Units (A.U.) and percentage positive for eGFP are reported as the geometric mean of three biological replicates) and cell viability (percentage of live single cells, reported as the arithmetic mean of three biological replicates); error bars ±SEM (standard error of the mean). **D**. Markers of pluripotency measured by nuclear intensity of OCT4 immunostaining via microscopy as shown in B and percentage of SSEA4 positive cells measured by flow cytometry. Data reported as the mean of three biological replicates; error bars ±SEM. **E**. Puromycin (Puro) or zeocin (Zeo) selection for 2 days enriches saRNA-expressing hiPSCs in 2D monolayers. Scale bar 200µm. **F**. Flow cytometry analysis of percentage of eGFP+ cells and mean eGFP expression at 4 dpt after 2 days of Puro or Zeo selection. Data reported as the mean of three biological replicates; error bars ±SEM. **G**. saRNA yields sustained, robust expression in 3D cultures. Cells are transfected in 2D hiPSC culture prior to seeding into suspension media for spheroid formation and culture over 4 days while measuring fluorescence intensity and spheroid size. Scale bar 300µm. **H**. Average eGFP fluorescence per unit area for cultures with and without Puro selection in suspension culture. N = 45 per distribution (5 spheroids per image and 3 images per well for 3 total wells). **I**. Maximum intensity projection of a representative spheroid from non-puromycin treated control and puromycin selected well imaged 5 days after seeding into 3D hiPSC spheroids. Scale bar 100µm. Statistical significance in **C**. and **D**. determined using one-way ANOVA with Bonferroni correction for multiple comparisons. Statistical significance in **F**. and **H**. determined using Student’s T test. ns (not significant) p>0.05; *p<0.05; **p<0.01; ***p<0.001; ****p<0.0001.

Here, we systematically test saRNA delivery of transgenes in various contexts, demonstrating their use to bridge the gap between rapid, integration-free delivery and durable expression in hiPSCs. Compared to transfection of modRNA and plasmid DNA, saRNAs efficiently induce sustained transgene expression in hiPSCs without affecting cell viability or pluripotency markers. A single transfection of saRNA encoding Ngn2 induces forward programming of hiPSCs to neurons at high efficiency within 6 days. We further demonstrate that delivery of saRNAs encoding functional reporters in hiPSCs can measure critical physiological readouts in hiPSC-derived cardiomyocytes, cardiac spheroids, and primary mouse hippocampal explants. Sparse-labeling allows measurement of cell physiology at single-cell resolution within mature 3D cardiac spheroids more than 30 days after reporter delivery. Collectively, our work demonstrates that saRNA-mediated delivery enables rapid, durable expression of transgenes, substantially increasing the speed and scalability of hiPSC engineering for a variety of applications.

## Results

### Self-amplifying RNA delivers efficient, durable transgene expression in hiPSCs

To characterize the potential of saRNAs to deliver sustained expression of transgenes to hiPSCs, we synthesized saRNA derived from components of the Venezuelan equine encephalitis (VEE) virus and compared expression to modRNA and transient transfection using a standard plasmid DNA (pDNA) vector (**Figure 1A**) [16]. The saRNA encodes the VEE nonstructural proteins (nsP1-4) responsible for amplification of the entire replicon and the transgenic cargo that is downstream of the subgenomic promoter (SGP). Using *in vitro* transcription, we first synthesized saRNA encoding an enhanced green fluorescent protein (eGFP) and a puromycin N-acetyltransferase (PuroR) selection cassette, separated by an internal ribosome entry site (IRES) downstream of the SGP (see Methods). We also synthesized modRNA and prepared pDNA encoding the same eGFP-IRES-PuroR cassette (**Figure 1A**), allowing us to directly compare across methods. hiPSCs were transfected with equal concentrations by mass of each construct. At 2 days post-transfection (dpt), we compared eGFP signal intensity using fluorescence microscopy (**Figure 1B**) and quantified expression by flow cytometry (**Figure 1C**). As expected, both modRNA and saRNA transfect a larger fraction of cells compared to pDNA (**Figure 1C**). We also observed that saRNA delivery leads to the highest transgene expression, whereby the mean level of eGFP expression using saRNA is 9.4-fold and 26.3-fold greater than modRNA and pDNA, respectively (**Figure 1B & C**).

Since delivery vectors and transgene expression can impact cellular physiology, we next measured hiPSC viability and pluripotency marker expression. At 2 days post transfection (dpt), cells retain high viability across all three delivery modalities, comparable to an empty transfection control (**Figure 1C**). We measured similar signal intensities of OCT4 immunofluorescence in the saRNA and modRNA conditions and observed a small reduction in pDNA transfected cells (**Figure 1D**). SSEA-4 levels were also similar across delivery methods as quantified by flow analysis (**Figure 1D).** Taken together, saRNA delivery achieves robust expression without affecting cell viability or expression of two key pluripotency markers.

Applications that require long-term expression such as lineage tracing and library screening may benefit from enriching for a population of cells expressing the transgene of interest. Thus, we generated saRNAs with either puromycin resistance (PuroR) or bleomycin resistance (BleoR) and performed antibiotic selection for 48 hours starting one day after transfection followed by media replacement without antibiotic (**Figure 1E**) [22], [23], [24]. After short-term selection, more than 90% of cells express eGFP at 4 dpt for both selection schemes. Puromycin treatment generates 1.5-fold higher levels of eGFP compared to zeocin treatment (**Figure 1F**). This difference may be due to the efficiency of the resistance enzyme or expression of the resistance cassette. Given the improved enrichment and expression levels of saRNA- expressing hiPSCs, we selected PuroR for use in all subsequent experiments.

Long-term experiments using hiPSCs to generate complex cell assemblies require durable expression of desired transgenes. Moreover, the transfection efficiency of 3D cell clusters is generally low, making delivery a challenge at this stage [25], [26]. We next tested whether delivery and antibiotic selection of saRNA in hiPSCs could overcome this limitation. saRNA-expressing hiPSCs were transfected and selected with puromycin for 7 days followed by generation of hiPSC spheroids [27] (**Figure 1G**). Puromycin selection in suspension culture for 3 days (starting the day after seeding) results in high quantities of eGFP-expressing hiPSCs across the cell population (days 2, 3, and 4 after seeding into suspension) (**Figure 1H & Figure S2D-E**). Potentially, modulating the cell seeding density and media volume can be used to tune spheroid diameter to accommodate for cell loss under puromycin selection [28], [29]. In the absence of selection, we observed sparse labeling of cells in spheroids (**Figure 1I**). The ability to achieve uniform and durable transgene expression or sparse labeling of cells provides flexibility for downstream applications [28], [29].

### saRNA-mediated delivery of Ngn2 efficiently differentiates hiPSCs to neurons

Forward programming of hiPSCs is a rapid, robust method to source rare or difficult-to-isolate patient- specific cells for disease modeling, drug discovery, and cell therapies [6]. For example, overexpression of the neuronal transcription factor Ngn2 rapidly differentiates hiPSCs to induced neurons (iNs) [30], [31]. Durable expression of Ngn2 improves neuronal purity [30]. As such, Ngn2-hiPSC lines typically use the Tet-ON 3G system, with a doxycycline-inducible Ngn2 transgene integrated into the genome [7], [32]. Due to the insertional mutagenesis risk associated with DNA-based delivery, integration-free methods of delivering TFs may reduce artifacts found in genome engineering and provide a greater safety profile for cell therapy applications [33]. While modRNA delivery of Ngn2 can drive hiPSC differentiation to iNs, the rapid decay of Ngn2 modRNA requires daily transfections for 10 days, substantially increasing cell stress and the cost and complexity of generating iNs.

To test the ability of saRNA-mediated Ngn2 expression for forward programming hiPSCs to induced neurons (iNs), we synthesized and delivered saRNA encoding Ngn2 and eGFP separated by a Porcine 2A (P2A) skipping peptide to hiPSCs, allowing us to correlate Ngn2 to eGFP expression (**Figure 2A**). After transfection, we cultured cells in conditions established for forward programming previously via a Tet-ON Ngn2 hiPSC line [34]. We tested differentiation with and without a 48-hour puromycin selection from days 1 to 3 (**Figure 2A**). By including the IRES-PuroR element on our construct, we can eliminate untransfected hiPSCs. A transient 48-hour puromycin treatment reduces the fraction of untransfected cells, yielding a higher proportion of iNs that form neurite projections (**Figure 2B**).

**Figure 2.**
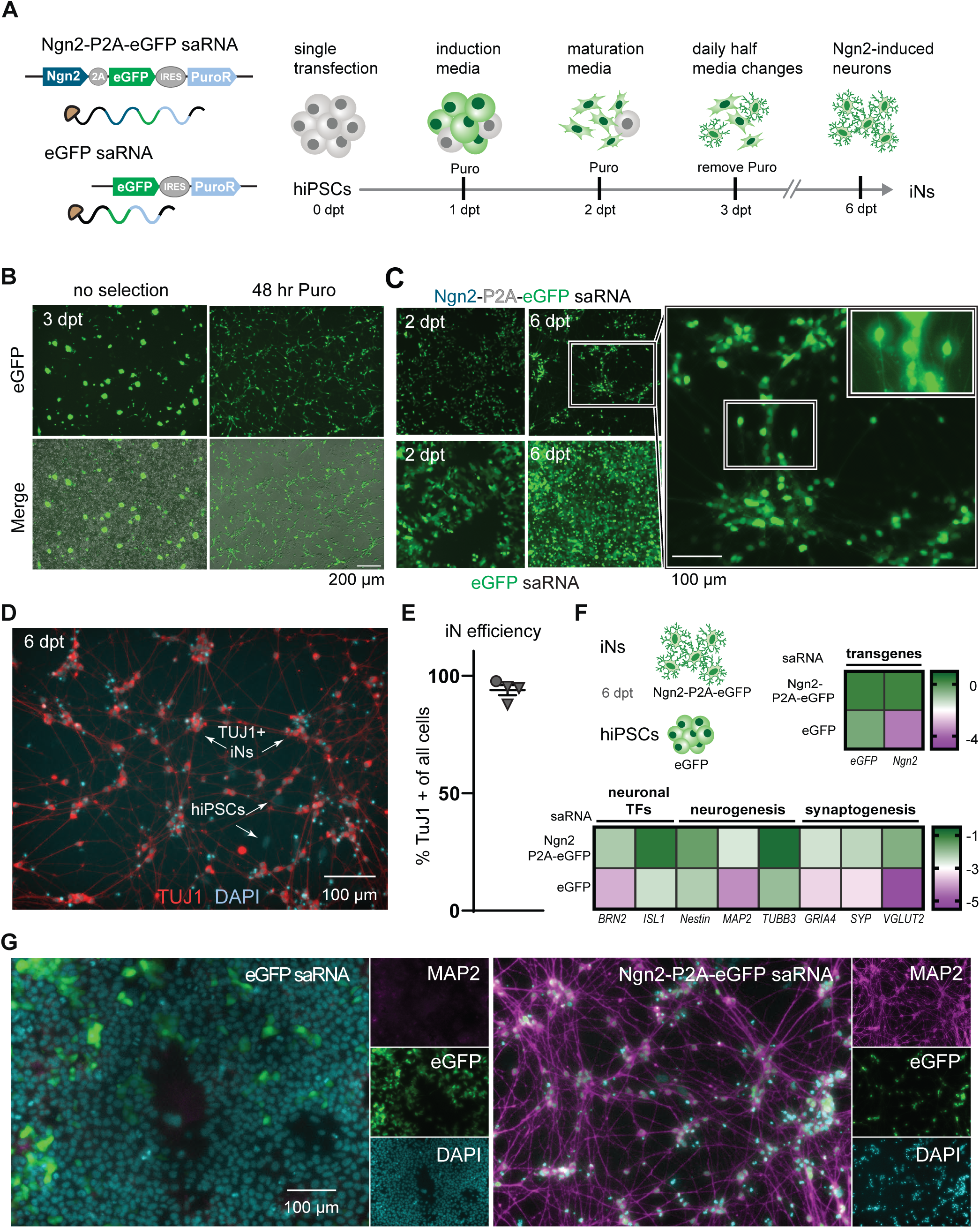
saRNA-based delivery of Ngn2 efficiently differentiates hiPSCs to induced neurons (iNs). **A**. Overview of saRNA-based forward programming of hiPSCs to iNs. **B**. Fluorescence microscopy images of hiPSCs transfected with Ngn2- P2A-eGFP saRNA taken at 3 dpt, with and without 48 hr Puromycin selection starting at 1 dpt. Scale bar 200 µm. **C**. Fluorescence microscopy images taken at 2 dpt and 6 dpt during conversion process of hiPSCs to iNs for hiPSCs transfected with either Ngn2-P2A-eGFP (top row) or eGFP saRNA control (bottom row). Scale bar 100 µm. **D**. Immunofluorescence images (TUJ1, red; DNA, cyan) of hiPSCs transfected with Ngn2-P2A-eGFP saRNA at fixed at 6 dpt, with white arrows indicating TUJ1+ iNs and TUJ1- hiPSCs (SCs). Scale bar 100 µm. **E**. Quantification of the percentage of TUJ1+ cells post differentiation at 6 dpt show high proportion of iNs. N = 2 independent differentiation experiments with 1 and 3 technical replicates (circle and triangle, respectively). **F**. Gene expression analysis of transgenes and neuronal markers, comparing the Ngn2-P2A-eGFP to eGFP saRNA control. Values represent the average of N = 3 independent differentiations, normalized to the RPL37A housekeeping gene and log10-transformed. **G**. Immunofluorescence images (MAP2, magenta; eGFP, green; DNA, cyan) of Ngn2-P2A-eGFP and eGFP saRNA control fixed at 6 dpt. Scale bar 100 µm.

With a single transfection of Ngn2-P2A-eGFP saRNA, we observed neurite projections from eGFP- expressing, puromycin-resistant cells and observed the emergence of neuronal clusters at 6 dpt (**Figure 2C**). The compact, spherical nuclei of iNs are readily distinguishable from the larger, elongated nuclei of hiPSCs (SCs) (**Figure 2D**). To measure differentiation efficiency, we fixed cells at 6 dpt and quantified the number of cells that are positive for TUJ1, by immunofluorescence, normalizing to the total number of nuclei. Delivery and selection yields cultures with more than 90% TUJ1 positive cells, indicating that Ngn2- P2A-eGFP saRNA induces highly efficient differentiation (**Figure 2E, Figure S4**). Delivery of eGFP saRNA does not yield TUJ1 positive cells despite culture in the same media conditions as the Ngn2-P2A-eGFP saRNA (**Figure S5**).

To measure activation of neuronal gene networks, we compared the expression of key neuronal genes in iNs and hiPSCs with eGFP saRNA. As expected, both Ngn2 and eGFP transgenes are expressed at 6 dpt as measured by RT-qPCR. Additionally, while eGFP is expressed in both saRNA transfected cell populations, only the Ngn2-P2A-eGFP saRNA transfected cells express Ngn2 (**Figure 2F**). Moreover, Ngn2-P2A-eGFP saRNA induces activation of neuronal genes including BRN2 and ISL1 compared to eGFP-transfected control (**Figure 2F**). Finally, as measured by immunofluorescence staining at day 6, iNs express MAP2, a marker associated with mature neurons [35] (**Figure 2G**). Collectively, these results indicate that saRNA-mediated delivery of Ngn2 drives highly efficient differentiation of hiPSCs into iNs that exhibit neuronal morphology, express neuron-associated genes, and are positive for mature neuronal markers with a single transfection of non-integrating genetic components.

### saRNA enables expression of functional reporters in primary and hiPSC-derived cells

Measuring real-time physiological changes in hiPSC-derived cells requires sustained expression of functional reporters. Silencing of genomically-integrated transgenes increases during differentiation, limiting the use of genetically-encoded tools for functional studies [9], [36], [37]. On the other hand, delivery of functional reporters using saRNA should resist silencing and allow durable expression. To test this idea, we constructed saRNAs encoding functional reporters representing diverse physiological readouts and localization to different subcellular compartments (**Figure 3A & Figure S6**). Specifically, we synthesized individual saRNAs carrying (i) ASAP5, a voltage reporter that localizes to the cell membrane (**Figure 3B**), (ii) jRCaMP1b, a red-shifted calcium reporter that localizes to the cytosol (**Figure 3C**), and (iii) HyPer7, a redox sensor targeted to mitochondria (**Figure S6**) [38], [39], [40].

**Figure 3.**
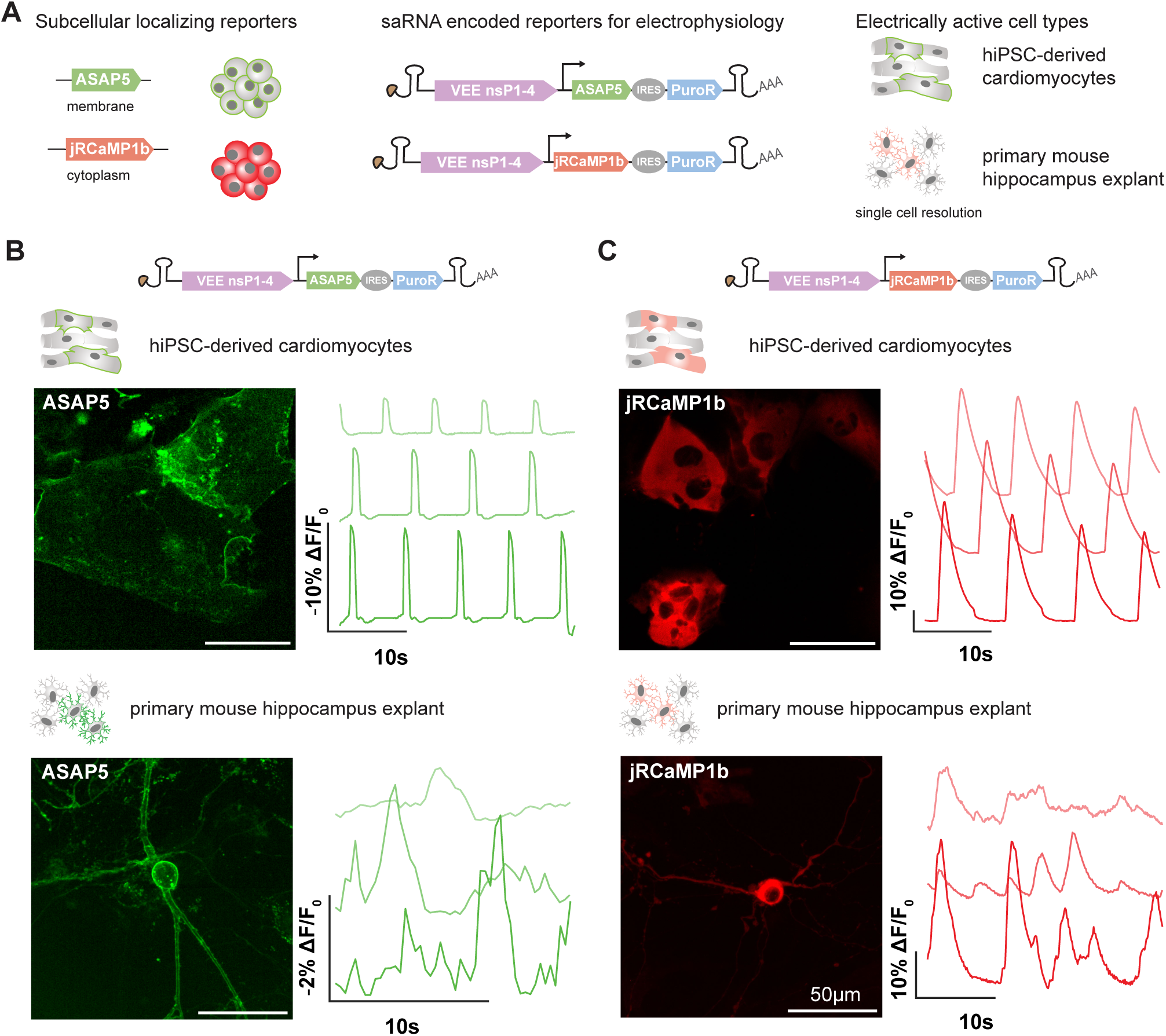
saRNA-encoded functional reporters properly localize in cells and report on voltage and calcium dynamics in electrically active cell types. **A**. Schematic of saRNA-mediated delivery of subcellular localizing fluorescent reporters to distinct cell types. For **B**-**E** N = 3 representative cells from a single transfection and all data collection performed at 2 dpt using a spinning disk confocal microscope (Yokogawa CSU-W1 confocal scanner unit on a Nikon Eclipse Ti microscope) equipped with a 40x 1.15 NA water immersion objective and Zyla PLUS 4.2 Megapixel camera. ASAP5 imaging (**B** & **C**) was performed using 488 nm excitation light and FITC emission filter and jRCaMP1b (**D** & **E**) imaging was performed using 561 nm excitation light and Cy3 emission filter. Fluorescence traces were normalized to a local baseline (F₀) calculated within a sliding window of W frames. For each frame, F₀ was defined as the Pth percentile of fluorescence values within that window, and the normalized signal is expressed as ΔF/F₀ = (|F − F₀|)/F₀. ASAP5 membrane localized voltage sensor in **B**. iPSC-derived cardiomyocytes (CMs) imaged at 10 Hz over 30 second recording period (normalization parameters: W = 25; P = 50) and **C**. primary mouse hippocampal explant culture at 3.33 Hz for 15 seconds (normalization parameters: W = 20; P = 90). Membrane depolarization events are reported as negative percent change from baseline fluorescence since ASAP5 exhibits inverse voltage sensitivity where brightness decreases during depolarization. jRCaMP1b cytosolic calcium reporter in **D**. iPSC-derived CMs imaged at 10 Hz for 30 seconds (normalization parameters: W = 100; P = 10) and **E**. primary mouse hippocampus explant cells imaged at 10 Hz for 30 seconds (normalization parameters: W = 100; P = 10). Scale bar 50 µm.

To assess sensor functionality, we tested saRNA-delivered reporters in cell types with robust electrical activity including hiPSC-derived cardiomyocytes (CMs) and primary neuronal cells that exhibit characteristic voltage and calcium dynamics. Measuring voltage changes using optical sensors remains challenging because of the fragile nature of the cell membrane, fast kinetics of voltage changes, and low expression of voltage reporters [41]. Thus, we selected ASAP5, a plasma membrane localized reporter, engineered for higher gain and faster onset enabling critical measurements and comparisons of electrical activity [39]. Next generation genetically encoded voltage indicators (GEVIs) such as ASAP5 offer a less invasive and higher throughput alternative to patch-clamp with enough spatiotemporal resolution to detect neuron firing and determine the shape of cardiac action potentials which differ between atrial, ventricular, and nodal cell types [42]. *In vivo*, ASAP5 shows reduced signal-to-noise and exhibits higher responsivity to action potentials and excitatory postsynaptic potentials compared to previously developed GEVIs. Delivery of saRNA encoding ASAP5 to primary mouse hippocampal cells, including neurons, and hiPSC-derived CMs shows ASAP5 signal localizes to the membrane (**Figures 3B & C**). As expected, ASAP5 traces appear as pulses of decreased fluorescence in response to membrane depolarization which showed stochastic activity in neurons and rhythmic patterns in CMs. In CMs, membrane depolarizations coincide with visible membrane contraction resulting in up to a 10% change in fluorescence over ∼1 second (**Figure 3B**). In addition to the characteristic action potential duration (APD) of cardiac muscle cells, ASAP5 traces exhibit a prolonged plateau phase and slower repolarization time which reflects the expected enrichment of ventricular CMs in hiPSC differentiation [43].

ASAP5 signal in mouse hippocampal culture was observed after 30 days *in vitro* and 2 days after saRNA transfection **(Figure 3C)**. At this timepoint, it is expected that primary hippocampal cultures will have formed network connections and exhibit spontaneous synaptic activity [44]. Within primary mouse hippocampus explant cultures, cells with characteristic neuronal morphology, including small cell bodies and long, branched processes, were visually identified and treated as putative neurons. Consistent with these expectations, we observe intermittent changes in reporter brightness which confirms spontaneous electrical activity in neurons as well as reporter responsiveness to depolarization events. For three representative cells recorded over 15 seconds, ASAP5 signal fluctuates by 1-5% over 1-2 seconds. These dynamics are consistent with subthreshold synaptic potentials as opposed to full action potentials [39]. Imaging modalities that achieve higher temporal resolution would further allow detection of the sub-millisecond action potentials associated with neurons, offering a potential alternative to more laborious patch clamp analysis.

To monitor calcium dynamics, we next transfected saRNA encoding jRCaMP1b, a genetically encoded calcium indicator (GECI) [38]. Like GEVIs, GECIs allow non-invasive measurement of cellular activity, but can be more readily observed on standard imaging setups because of their slower rate of change and higher expression throughout the cytoplasm (as opposed to the cell membrane). Delivery of saRNAs encoding jRCaMP1b to hiPSC-derived CMs and primary mouse hippocampus explants enabled measurement of calcium transients in single cells (**Figures 3D & E**). As with ASAP5, cardiomyocytes display characteristic rhythmic patterns of calcium signaling (**Figure 3D**) and neurons show stochastic activity across cells like those reported for GECIs (**Figure 3E**) [45]. Across three representative cells, jRCaMP1b signal fluctuates by a maximum of 8-15% over an average of 6.67 seconds in CMs with a faster increase in fluorescence lasting 1.13 seconds and longer return to baseline in 5.53 seconds. This is consistent with spontaneous calcium transients in immature hiPSC-derived CMs recorded at room temperature, where reduced sarcoplasmic reticulum function and slower ion channel kinetics prolong rise and decay times [46], [47], [48]. The observed amplitudes fall within the range expected for red-shifted genetically encoded calcium indicators [38]. In putative mouse neurons, jRCaMP1b fluorescence increased by up to 31%, and a single peak lasted at most 5.86 seconds. These amplitudes and kinetics are consistent with previous reports of jRCaMP1b in cultured neurons, where transient durations ranged from 1-10 seconds [38], [49]. The relatively slow decay of the signal suggests that these neurons exhibit prolonged intracellular calcium elevations, potentially reflecting sustained spontaneous activity under these experimental conditions. We also demonstrate saRNA can deliver mitochondrially targeted HyPer7, a genetically encoded fluorescent H2O2 sensor that can be read out in diverse cell types (**Figure S6**). Thus, saRNA offers a versatile platform for rapid, non-integrating delivery and durable measurement of functional reporters in primary and differentiated cell types using conventional fluorescence microscopy.

### saRNA delivery of reporters enables functional readouts in hiPSC-derived 3D cardiac spheroids

To examine the potential of saRNA-delivered fluorescent reporters in monitoring hiPSC differentiation in 3D systems, we tested the durability of saRNA-delivered transgenes during hiPSC differentiation to cardiac spheroids. Remarkably, we demonstrate that saRNA-mediated transgene expression persists throughout differentiation for over 30 days (**Figure 4A**). We transfected iPSCs with saRNA in 2D adherent culture and selected with puromycin for 5-7 days prior to cardiac differentiation. saRNA-enriched hiPSCs were seeded into spheroids followed by cardiac differentiation using standard protocols (see Methods). We observed beating between days 6 and 7 of differentiation in saRNA-eGFP labeled cardiac spheroids. Immunofluorescent staining shows robust expression of CM markers Nkx2.5 and cTnT by day 14 of differentiation, including in cells that express eGFP (**Figure 4B**). saRNA-treated CMs also exhibit a clear sarcomere banding pattern by cTnT immunofluorescence staining We next tested a 24-hour puromycin selection step to further enrich cells expressing saRNA once seeded into suspension. Across replicates, puromycin selection increased the fraction of cells labeled at day 14 (**Figure 4C**). To maximize expression of the reporter in cTnT positive cells at day 30, we included this enrichment step for experiments continued to day 30 (**Figure 4G**), but omitted selection for day 14 measurements (**Figures 4D & E**). The brief additional selection does not affect the generation of robustly beating eGFP-labeled spheroids that are positive for Nkx2.5 and cTnT (**Figures S8**).

**Figure 4.**
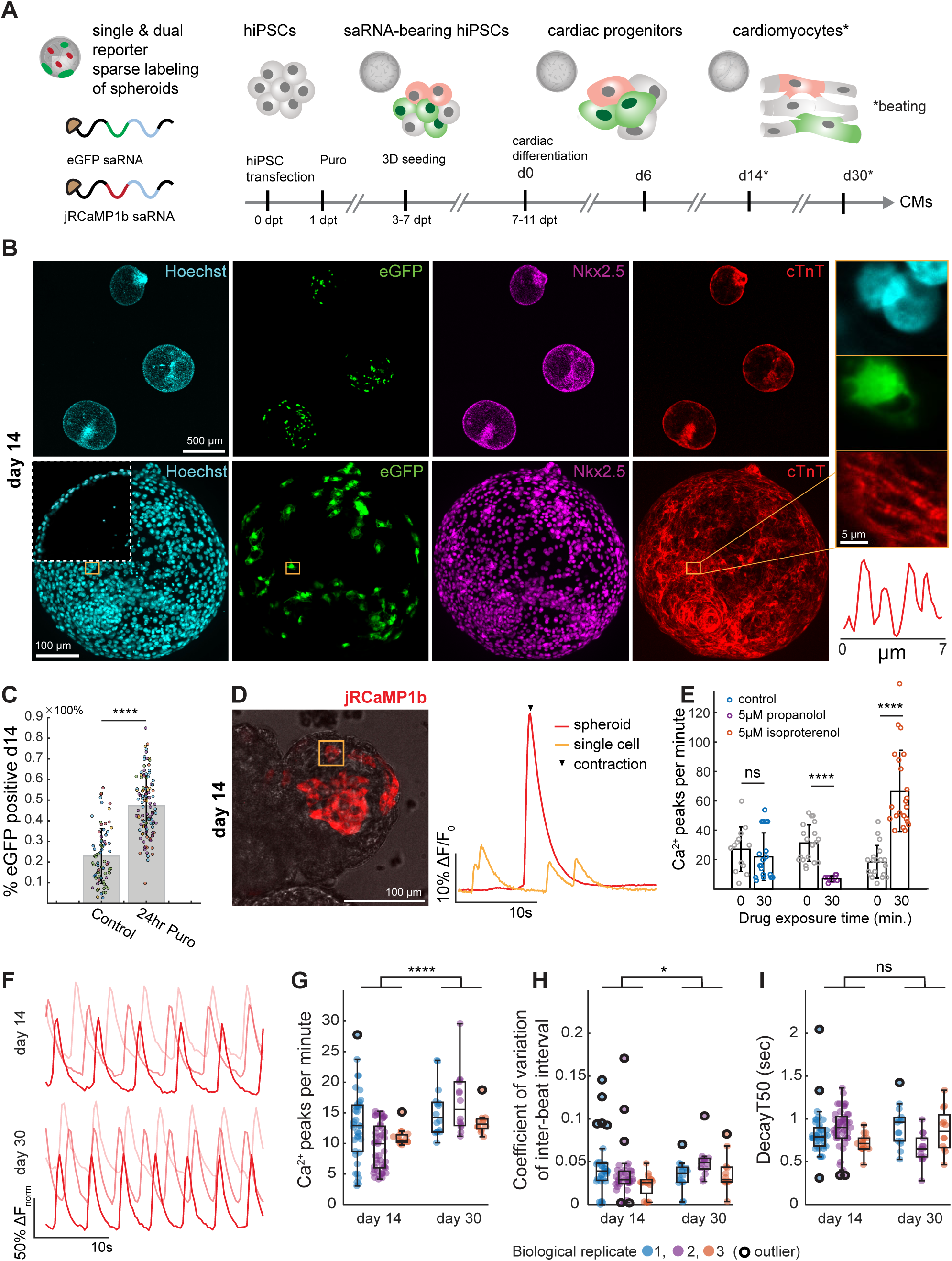
saRNA-based fluorescent reporters enable long-term functional readout of calcium dynamics in 3D cardiac spheroids. **A**. Overview of saRNA-based reporter delivery to hiPSCs to cardiac spheroid differentiation protocol. **B**. Maximum projections of immunofluorescent images (Hoechst, cyan; eGFP, green; Nkx2.5, magenta; cTnT; red) of a representative cardiac spheroid expressing saRNA-eGFP fixed at day 14 of differentiation. Top row: 4x magnification, Scale bar 500 µm; Bottom row: 20x magnification, Scale bar 100 µm. Inset with dotted white line shows a single z-slice of Hoechst channel demonstrating chamber formation. Inset with orange border shows single z-slice at 40x magnification showing banded sarcomere formation in eGFP positive cell, Scale bar 5 µm. **C**. Labeling percentage for day 14 cardiac spheroids expressing saRNA-eGFP with and without an initial 24-hour puromycin (Puro) selection step after seeding into 3D suspension culture. Each point represents a single spheroid, and each distribution contains two independent differentiations. (Control: N = 77 from two differentiations with 3 wells each, 24hr Puro; N = 115 from two differentiations with 2 wells and 3 wells each). Data are reported as the mean and error bars represent standard deviation. p = 8.25x10^-25^ by T-test. **D**. (left) Cardiac spheroid at day 14 of differentiation expressing saRNA-jRCaMP1b. Representative overlay of brightfield and maximum projection of z-stack at 561 nm illumination taken at 40x on day 14 of differentiation. (right) Accompanying 30 second recording of average change in jRCaMP1b signal measured in a single cell (orange line) at sub-contraction level and across the entire cardiac spheroid (red line) during a contraction, noted with black arrow. Normalized by minimum fluorescence value across recording period. Scale bar 100 µm. **E**. Calcium peaks per minute as measured by 30 second recordings of jRCaMP1b signal at 4x magnification with 300 ms exposure time (9 fps) before and after 30-minute incubation with 5 µM propranolol or 5 µM isoproterenol compared to control media (from left to right N = 13, 20, 20, 14, 19, 23 beating cardiac spheroids). Data reported as the mean of all observations; error bars represent standard deviation. p = 0.3704, 3.354x10^-8^, 1.022x10^-8^ by T-test. **F**. Representative traces of jRCaMP1b taken at day 14 and day 30 of three independents differentiations. One representative trace selected per differentiation. Normalized between 0 and 1 to compare temporal parameters. **G**. Calcium peaks per minute, **H**. coefficient of variation of inter-beat interval, **I**. time to decay to 50% (Decay T50) of peak brightness for three independent differentiations compared at day 14 and day 30 where N = 38, 49, 16 for day 14 and N= 16, 14, 12 for day 30 across 3 wells each (day 14; N = 16 represents one well). For **G**-**I** data are represented by a boxplot showing the median (line within box), interquartile range/IQR (box), and data points outside of 1.5 x IQR as outliers. Whiskers extend to the most extreme points within 1.5 x IQR. p = 7.67x10^-7^, 0.017, and 0.3771 for **G**-**I** respectively by Mann-Whitney U test. ns (not significant) p>0.05; *p<0.05; **p<0.01; ***p<0.001; ****p<0.0001.

Having demonstrated maintenance of saRNA-eGFP expressing spheroids throughout a 30-day 3D differentiation, we next tested our ability to measure a dynamic physiological reporter in this system. Sparse labeling is particularly valuable for dynamic imaging, such as calcium or voltage signals, where overlapping fluorescence from neighboring cells can obscure spatial resolution [50]. Calcium fluxes directly reflect excitation–contraction coupling and electrophysiological maturity and is a gold-standard functional assay for hiPSC-derived CMs in disease modeling and drug screening contexts [51], [52]. Thus, we generated sparsely labeled jRCaMP1b expressing cardiac spheroids and observed calcium transients during differentiation progression. At day 14, we measured both small, non-contractile fluctuations in single cells and large, contraction-coupled calcium transients across the entire spheroid plotted as normalized fluorescence intensity (ΔF/F0) over time (**Figure 4D**). Cardiac spheroids expressing both eGFP and jRCaMP1b can also be generated by seeding equal numbers of eGFP-transfected and jRCaMP1b- transfected hiPSCs prior to differentiation making this a flexible method for combining multiple labels without burdening individual cells with more than one transgene (**Figure S8**). As proof of concept, we next measured calcium dynamics using jRCaMP1b expressing CMs in response to well characterized drugs that disrupt cardiomyocyte function. We added propranolol or isoproterenol, that block or increase calcium influx, respectively, to day 14 cardiac spheroids for 30 minutes and measured fluorescence intensity over time. From a single field of view with low magnification, we could segment and extract up to 20 independent calcium recordings per condition to rapidly assess changes in beat rate due to drug treatment. As expected, exposure to propranolol decreased calcium peak number by 77.3% and treatment with isoproterenol increased the calcium peak number by 257%, while the control group shows no statistically significant change (**Figure 4E & Figure S9**).

The ability to maintain reporter expression for more than 30 days using saRNA delivery enables measurement of calcium dynamics in CMs throughout differentiation and maturation (**Figure 4F-I and Figure S8**). To track calcium signaling over a longer observational period, we include 24-hour puromycin selection upon seeding into suspension to increase reporter signal at day 30. Consistent with CM maturation, we found that beat rate increases from day 14 to 30 across three independent differentiations. Moreover, the coefficient of variation (CV) of the interbeat interval is slightly higher on average at day 30 compared to day 14. The narrower distribution at day 30 indicates that there are fewer spheroids with irregular beating patterns observed across the population (**Figure 4H**). As cardiomyocytes mature, they develop faster calcium decay kinetics, a key hallmark of functional maturation and efficient excitation-contraction coupling [53]. Despite a higher average number of calcium peaks per minute, we do not observe statistically significant differences in calcium decay kinetics at day 30 compared to day 14 (**Figure 4I**). These data are consistent with the known lack of functional maturity of hiPSC-derived CMs indicating the need for assessing calcium dynamics beyond beat frequency. Although we initially focused on parameters that prioritize experimental throughput to support the idea that saRNA can scale to screening applications, we found that sensor brightness at day 30 is sufficient to capture calcium transients at a 50 Hz frame rate enabling faster kinetic measurements, including rise time (**Figure S8**). Moreover, saRNA- GECIs optimized for fast transient calcium imaging can be used if higher temporal resolution or speed is preferred over the longer-lasting spectral properties of jRCaMP1b. Taken together, we demonstrate that saRNA-encoded fluorescence reporters can be used to track dynamic signals over long observational periods with a single non-integrating transfection, greatly expanding the toolkit for non-invasive imaging of biological processes for multiple applications that require extended culture time.

## Discussion

Here, we systematically test saRNA delivery to hiPSCs and primary cells as a flexible tool for durable expression of transgenes in multiple applications. saRNAs generate high protein levels in hiPSCs without compromising their viability, pluripotency, or ability to differentiate. We also show that a single transfection of saRNA encoding *Ngn2* to hiPSCs leads to high efficiency differentiation to neurons. Moreover, we demonstrate that saRNAs can deliver genetically encoded voltage or calcium indicators to electrically active cell types such as neurons and cardiomyocytes. Finally, we show that saRNA delivery of jRCaMP1b to hiPSCs allows long-term monitoring of calcium dynamics to assess maturity and response to drug perturbations in cardiac spheroids. Thus, saRNA offers a rapid, efficient method to forward program hiPSCs to neurons and to generate footprint-free fluorescent reporter lines. Together our data support that saRNA can be broadly used as a highly scalable tool for functional manipulation and observation of hiPSC and hiPSC-derived cells.

Current methods to introduce genetic payloads to hiPSCs are subject to a major tradeoff between duration of expression and risk from genomic integration. Non-integrating tools including Sendai virus, episomal vectors, synthetic mRNA, plasmid DNA, protein transduction, and AAV pose little risk for unintended genomic integration. However transient expression from these tools is often insufficient for observing differentiation processes or longer-term monitoring of disease states [54]. Integrating tools like lentivirus, retrovirus, CRISPR/Cas9, and transposons are often used to generate stable cell lines, but these methods pose a higher risk of insertional mutagenesis, off-target effects, and genetic drift [55], [56]. Leveraging saRNA to facilitate long-term transgene expression obviates the need for the cumbersome and slow pipelines of cellular engineering, enabling more rapid exploration of biological phenomena. The ease with which saRNA tools achieve high efficiency, long-term transgene delivery serves to democratize stem cell engineering and enable more labs to incorporate biosensors in their measurements of hiPSCs and hiPSC-derived cell types. This will accelerate the rate at which candidate factors and biosensors can be tested across tissue systems, driving the development of next-generation therapies and advancing new discoveries [57].

We demonstrate the applicability of saRNA to forward program hiPSCs to induced neurons. *Ngn2*-driven differentiation of hiPSCs yields neurons that can be used for drug screening and cell therapy. Previous work established that forward programming to neurons requires durable expression of Ngn2 [30]. As such, Ngn2-hiPSC lines typically use the Tet-ON 3G system, with a doxycycline-inducible Ngn2 transgene integrated into the genome along with a constitutively expressed transactivator [7], [32]. Depending on the syntax of the Tet-ON elements, a single site-specific integrated copy of tet-inducible Ngn2 can efficiently differentiate hiPSCs to iNs [58]. However, harnessing the power of diverse hiPSC lines requires scalable methods of engineering neurons, underscoring the value of fast, robust tools. Importantly, as a non-integrating vector with FDA approval for administration to patients [59], saRNA may offer improved scalability and safety for translating hiPSC-derived neurons as cell therapies. The simple, rapid production of hiPSC-derived induced neurons at clinically relevant purities via Ngn2 saRNA provides allows scaling the potential of disease-modeling and drug screening with hiPSC-derived cells. Combining saRNA *Ngn2*- driven forward programming with specific small molecules and growth factors may further advance disease modeling by enabling the scalable generation of neuronal subtypes [60], [61], [62].

3D spheroids and organoids have emerged as powerful models for studying human development, disease, and drug responses, offering a more physiologically relevant alternative to traditional 2D cultures. Their 3D architecture and multicellular complexity replicate key features of native tissues, but these same properties create challenges for gene delivery. This limitation is especially true for non-integrating, transient transfection methods like lipid-based reagents or electroporation, which fail to deliver payloads within dense, differentiated structures. Thus, introducing genetic tools, such as fluorescent biosensors, earlier in the workflow by manipulating hiPSCs in 2D adherent culture could overcome this limitation. We demonstrate that saRNA-based delivery achieves sparse sensor labeling with robust and enduring signal in 3D cardiac spheroids with a single transfection prior to differentiation. Since saRNA delivery is highly scalable, its implementation to deliver reporters across many cell lines offers a key advantage for the identification of genotype-specific effects of drug compounds, thereby improving genotype-based stratification of candidates for clinical trials. Moreover, the durable expression of fluorescent reporters avoids repeated dosing while enabling dynamic measurements for long-term safety studies, a feature lacking in traditional drug screening modalities [63]. Though we initially selected cardiac spheroids as a testbed system to demonstrate the utility of saRNA to construct footprint-free optical reporter lines, we expect this approach to be applicable across multiple tissue types, further augmenting the potential of saRNA as an impactful addition to a researcher’s toolbox [59].

Beyond our demonstrations in forward programming and measuring functional reporters, there remain opportunities to tailor the design of saRNAs for their broadest application. High expression of transgenes supported our testbed applications of Ngn2-based differentiation to neurons and calcium tracing in cardiac spheroids. However, some applications may require precise or controllable levels of protein levels. For example, transient expression of the pioneer transcription factor ETV2 in hiPSCs produces more mature endothelial cells compared to sustained overexpression [64]. Moreover, overexpression of some genetic sensors may induce proteotoxic stress, metabolic burden, or other off-target physiological effects [65]. Synthetic biology approaches to control saRNA transgene expression can support tuning of expression.

For example, subgenomic promoter tuning [66] can offer pre-translational control of transgene expression, and fusing TMP-responsive degradation domains to the alphaviral non-structural proteins or the transgene product offers post-translational control [67]. Combining evolving control mechanisms could further tune output of transgene expression by calibrating initial expression level and offering inducible degradation of replication machinery and/or gene product. In theory, multiple transcription factors or combinations of transcription factors and reporters could be encoded on a single saRNA. However, the design rules for larger polycistronic cassettes are not yet defined. Both the exact sequence of the transgene, their connecting sequences (e.g., IRESs and self-cleaving peptides), and the length of saRNA may affect transfection and expression. However, the modularity of saRNA-encoded payloads should enable rapid and iterative design testing across combinations of genetic components to optimize performance. Lastly, cytotoxicity can differ based on the specific saRNA vector (e.g., New World versus Old World alphaviruses) used and can also be mitigated using immune-evasive saRNA that intrinsically suppresses innate immune response pathways or by complete substitution with modified nucleotides [68],[69].

Collectively, our study shows that saRNA is a fast, simple, robust, and non-integrating method for transgene delivery offering a more scalable solution for engineering hiPSCs. With broader adoption of saRNA-based tools in this context, we anticipate accelerated innovations and discoveries across developmental biology, disease research, and drug screening applications.

## Lead contact

Requests for further information and resources should be directed to and will be fulfilled by the lead contact Laurie Boyer, PhD and Kate E. Galloway, PhD.

## Materials availability

All plasmids will be made available on Addgene.

## Data and code availability

Custom scripts and analysis code supporting the findings of this study will be made publicly available on GitHub upon publication. Any additional information required to reanalyze the data reported in this paper is available from the lead contact upon reasonable request.

## Acknowledgements

Research reported in this manuscript was supported by the National Institute of General Medical Sciences of the National Institutes of Health under Health under award number R35-GM143033, by the National Science Foundation under the NSF-CAREER under award number 2339986, by the Air Force Research Laboratory MURI under award FA9550-22-1-0316, and funding from Institute for Collaborative Biotechnologies under cooperative agreement W911NF-19-2-0026 (K.EG). This work was also supported by the National Institute of Heart Lung and Blood R01 HL140471 (L.A.B.). We thank MIT M-CELS seed funding for supporting this work (K.E.G. and L.A.B.). E.S.B. acknowledges funding from HHMI, Lisa Yang, and NIH R01 AG087374.

We thank the Koch Institute’s RA. Swanson (1969) Biotechnology Center (National Cancer Institute Grant P30-CA14051) for technical support, specifically the Microscopy Core and the BioMicro Center Core facilities. We thank Nat Wang, Leah Borden, and Dandan Yang for experimental help. We thank Dandan Yang, Nat Wang, Brittany Lende-Dorn, and Maria Castellanos for feedback on the manuscript.

## Author contributions

C.D.S., D.S.P., K.E.G., and L.A.B. conceived and outlined the project. C.D.S., D.S.P., and N.L.M. conducted flow cytometry and immunofluorescence comparison of saRNA, modRNA, and pDNA. D.S.P. performed the molecular cloning, synthesized modRNA and saRNA, and analyzed data. D.S.P., M.E.E, and A.B.A conducted neuronal differentiations and immunostainings. C.D.S. performed transfection, imaging, and analysis of voltage and calcium reporters. E.C.K. prepared primary neural cultures and advised transfection experiments. C.D.S. and C.G.B. performed cardiac spheroid differentiations, imaging, and analysis. C.D.S., D.S.P., L.A.B., and K.E.G. wrote and edited the paper with input from all authors. L.A.B., K.E.G., and E.S.B. secured funding and supervised the project.

## MATERIALS AND METHODS DETAILS

Reagents and resources table (in separate document)

### Experimental model and study participant details

All procedures involving animals at the Massachusetts Institute of Technology were conducted in accordance with the United States National Institutes of Health Guide for the Care and Use of Laboratory Animals and approved by the Massachusetts Institute of Technology Committee on Animal Care and Biosafety Committee.

iPS11 cells (Alstem, iPS11) were maintained on 24-well plates coated with 15 µL CellAdhere™ Laminin-521 (STEMCELL Technologies, 200-011), media changed daily with 500 µL Gibco™ Stemflex™ (ThermoFisher Scientific, A3349401) containing 100 U/mL Penicillin-Streptomycin (ThermoFIsher Scientific, 15140122) and incubated in a humidified incubator at 37°C with 5% CO2. Cells were passaged at a split ratio of 1:10 twice a week when ∼80% confluent using 300 µL Gentle Cell Dissociation Reagent™ (STEMCELL Technologies, 100-1077) according to the manufacturer’s instructions and replated with 1:200 RevitaCell™ Supplement (ThermoFisher Scientific, A2644501). iPS11 are used in Figures 1B-F and 2B-G.

37-1189 cells (generously provided by University of Pennsylvania under MTA agreement, PENN0141-37- 3) were maintained on 6-well plates coated with hESC qualified Matrigel (Corning, 354277) diluted in 500 mL DMEM/F12 media (Gibco, 11320033), media changed every other day with 2 mL Gibco™ Stemflex™ (ThermoFisher Scientific, A3349401) and incubated in a humidified incubator at 37°C with 5% CO2. Cells were passaged at a split ratio between 1:3 and 1:6 when <60% confluent using 1 mL ReLeSR^TM^ (STEMCELL Technologies, 100-0483) according to manufacturer’s instructions. 37-1189 cells are used in Figures 1G-H and 4B-I.

hiPSC-derived CMs were obtained by differentiation of 37-1189 hiPSCs [72], [73]. hiPSCs were maintained on Matrigel-coated plates (Corning, 354277) in mTeSR Plus medium (STEMCELL Technologies, 100- 0276), with medium changes every two days. Cells were passaged using ReLeSR (STEMCELL Technologies, 100-0483) according to manufacturer instructions. For differentiation, cells were seeded onto Matrigel (Corning, 354277) in a 12-well plate at a concentration of 100,000 cells per well, allowed to grow for 3 days, and then treated with 12 µM CHIR-99021 (Selleck Chemicals, S2924) for 24 hours in RPMI 1640 (ThermoFisher Scientific, 11875093) with B27 supplement, minus insulin (ThermoFisher Scientific, A1895601). Exactly 24 hours after removal of CHIR-99021, the cells were treated with 5 µM of the Wnt inhibitor IWP-2 (Fisher Scientific, 35-331-0) and incubated for another 48 hours. RPMI/B27-insulin media was changed every 2 days. 7 days after treatment with CHIR-99021, the media was switched to RPMI with B27 supplement, serum free (ThermoFisher Scientific, 17504044). To select for cardiomyocytes, cells were cultured for 3 days in RPMI 1640 without glucose (ThermoFisher Scientific, 11879020) supplemented with 213 µg/mL L-ascorbic acid 2-phosphate (Sigma-Aldrich, TA9H9876E755), 500ug/mL human albumin (Sigma Aldrich, A9731), and 5mM sodium L-lactate (Sigma Aldrich, 71718). Cardiomyocytes were replated using the STEMdiff™ Cardiomyocyte Dissociation Kit (StemCell Technologies 05025) and plated onto a Matrigel-coated plate for imaging at day 14. hiPSC CMs were used in Figure 3B & C.

Primary mouse hippocampal explant cultures were prepared from postnatal day 0 or 1 Swiss Webster mice (Taconic) (both male and female mice were used) as previously described (Klapoetke et al., 2014 [74]). Briefly, dissected hippocampal tissue was digested with 100 units of papain (Worthington Biochem) for 6- 8 min, and the digestion was stopped with ovomucoid trypsin inhibitor (Worthington Biochem). Cells were seeded in 100 μl neuron culture medium containing Minimum Essential Medium (MEM, no glutamine, no phenol red; Gibco), D-glucose (25 mM, Sigma), holo-Transferrin bovine (100 µg mL^−1^, Sigma), HEPES (10 mM, Sigma), glutaGRO (2 mM, Corning), insulin (25 µg ml^−1^, Sigma), B27 supplement (1×, Gibco), and heat-inactivated fetal bovine serum (10% in volume, Corning), with final pH adjusted to 7.3–7.4 using NaOH. After cell adhesion, additional plating medium was added. AraC (2 µM, Sigma) medium was added at 1 day *in vitro* (DIV 1), when glial density was 50%–70% of confluence. Neurons were grown at 37°C and 5% CO2 in a humidified atmosphere in a neuron incubator, with 2 mL total medium volume in each well of the 24-well plate. Primary mouse hippocampal explant cultures are used in Figure 3B & C.

### Molecular cloning

All self-amplifying RNA IVT templates were derived from T7-VEE-GFP (Addgene Plasmid #58977). To streamline saRNA production, T7-VEE-eGFP-IRES-PuroR-polyA (Addgene Plasmid #246033), which encodes a polyA track downstream of the 3’ conserved sequence element was constructed. This enables T7 RNA polymerase to append a polyA tail on transcribed saRNAs within the IVT reaction, eliminating the need to carry out an additional polyadenylation reaction required for saRNAs derived from the original vector. To achieve this, a 4 nmol DNA ultramer oligo (Integrated DNA Technologies), polyA-encoded, which contained a segmented polyA (poly(A)2 × 60_G) previously reported to attenuate in vivo truncation in E. coli [75] flanked with Gibson overhangs was ordered. T7-VEE GFP was digested with Sph-HF^®^ and MluI-HF^®^, which corresponded to the unique restriction sites closest to the desired placement of a polyA track. As the digestion of T7-VEE GFP with Sph-HF^®^ excised the C-terminus of PuroR and the 3’ conserved sequence element, the corresponding sequence from T7-VEE GFP was PCR amplified with primers DP- 369 and DP-370. These and all subsequent digested vectors and PCR-generated DNA fragments were isolated via gel electrophoresis, extracted with Agarose Dissolving Buffer (Zymo Research, D4001-1-100), and purified with the Monarch PCR and DNA Cleanup Kit (NEB). 40 ng of digested backbone, 40 ng PCR product, and 4 ng of polyA-encoded oligo were assembled using HiFi reaction mixture (NEB). The reaction was transformed to NEB stable and incubated overnight on LB+Ampicillin Agar plates. Individual colonies were cultured in 5 mL liquid LB+Ampicillin, with overnight cultures miniprepped using the Monarch miniprep kit. All incubation steps were conducted at 30° C to minimize truncation of the polyA track.

All saRNA IVT templates were assembled via HiFi assembly following an analogous procedure. The eGFP- IRES-PuroR cassette from T7-VEE-eGFP-IRES-PuroR-polyA was excised by digestion with AscI and XbaI. The digested backbone was treated with shrimp alkaline phosphatase to attenuate background colonies. Genes of interest were PCR amplified with 20-30 bp long Gibson-compatible overhangs with PrimeSTAR® HS DNA Polymerase (Takara). BleoR was cloned from pIVT-Flp-2A-BleoR (Addgene Plasmid #232752), jRCaMP1b was cloned from pAAV.Syn.NES-jRCaMP1b.WPRE.SV40 (Addgene Plasmid #100851), ASAP5 was cloned from pAAV-hsyn-ASAP5-WPRE (Addgene Plasmid # 225709), HyPer7 was cloned from HyPer7.2DAAO-mito (Addgene Plasmid #168304), GINKO1 was cloned from pcDNA3.1-GINKO1 (Addgene Plasmid #113112) and Ngn2-P2A was cloned from pMXs-NIL (Addgene Plasmid # 233192).

The template for eGFP-IRES-PuroR modRNA, pIVT-eGFP-IRES-PuroR-polyA (Addgene Plasmid #246041) and eGFP-IRES-PuroR plasmid DNA expression vector, pSHIP-CAG.d-eGFP-IRES-PuroR-bGH (Addgene Plasmid #246040) were both constructed via BsaI Golden Gate following the Galloway Lab’s version of MoClo, as previously reported [76] . Briefly, the eGFP-IRES-PuroR cassette was PCR amplified with compatible pPV2 BsaI overhangs. For pIVT-eGFP-IRES-PuroR-polyA, the PCR fragment was assembled into a modRNA IVT backbone plasmid containing beta globin 5’ and 3’ UTRs and a T7 promoter compatible with the CleanCap AG reagent. For pSHIP-CAG.d-eGFP-IRES-PuroR-bGH, the PCR fragment was assembled into a transcriptional unit containing a domesticated CAG promoter (CAG.d) and the bovine growth hormone (bGH) polyadenylation signal. Golden gate reactions were transformed as described above, substituting Ampicillin with Kanamycin.

All plasmids were sequence verified with Plasmidsaurus Whole Plasmid Sequencing and will be made available on Addgene.

### *In vitro* transcription

Plasmids encoding the desired saRNAs were linearized for IVT with MluI-HF^®^ (New England Biolabs, R3198S) and purified using the Monarch PCR and DNA Cleanup Kit (New England Biolabs, T1030). 500 ng of purified linearized product served as template in a 20 µL IVT reaction using the HiScribe T7 High Yield RNA Synthesis Kit (New England Biolabs, E2040), co-transcriptionally capping with CleanCap Reagent AU (TriLink Biotechnologies, N-7114-1) for saRNA or CleanCap Reagent AG (TriLink Biotechnologies, N-7113-5) for modRNA. 1 µL of Pyrophosphatase, Inorganic (E. coli) (New Englad Biolabs, M0361S) was added to each IVT to enhance yield. IVT reactions were incubated at 37°C for 6 hours, at which point reactions were diluted to 50 µL, treated with 2 µL DNase I (New England Biolabs, M0303), and incubated at 37°C for 30 min to degrade the IVT template DNA. Synthesized saRNA was column purified and eluted with 60 µL water using the 50 µg Monarch RNA Cleanup Kit (New England Biolabs, T2040). A small sample was nanodropped and run on a native denaturing gel to determine saRNA concentration and verify full-length product. The saRNA was dispensed in aliquots and stored at −80°C.

### Transfection of hiPSCs with saRNA

Transfection experiments were conducted as described elsewhere [58]. Briefly, hiPSCs were plated the day prior at 100,000 cells per 24-well to achieve ∼30% confluency on the day of transfection. 500 ng of saRNA was diluted in Opti-MEM™ (ThermoFisher Scientific, 31985070) to a volume of 25 µL. In parallel, either 1 µL of Lipofectamine™ Stem Transfection Reagent (ThermoFisher Scientific, STEM00015; for Figure 1 comparisons of plasmid DNA, modRNA, and saRNA) or 1 µL of Lipofectamine™ MessengerMAX Transfection Reagent (ThermoFisher Scientific, LMRNA008; for all other experiments involving exclusively modRNA and/or saRNA) was added to 24 µL Opti-MEM™. For the Stem Transfection Reagent, the diluted saRNA was added immediately to the Lipofectamine™ mixture, mixed well, and incubated for 10 minutes. For the MessengerMax Transfection Reagent, the Lipofectamine™ mixture was preincubated for 15 minutes before adding the diluted saRNA, and the saRNA/Lipofectamine™ mixture was incubated for an additional 5 minutes. During the incubation periods, cells were optionally washed once with PBS, and media was replaced with 500 µL of Opti-MEM™. After the final incubation step, the saRNA/LNP complexes were added drop wise to the well, and the plate was gently rocked and returned to the incubator. After 4 hours, 500 µL of StemFlex media was added to each transfected well. A complete media change was performed on transfected wells the following day.

### Quantification of transfection efficiency, cell viability, and pluripotency markers

iPS11 cells were transfected as above with 500 ng of either pDNA, modRNA, saRNA using Lipofectamine™ Stem Transfection Reagent. Two days post-transfection, cells were either fixed and stained for OCT4 or dissociated with TrypLE Express^TM^ for flow cytometry assessment of viability and SSEA-4 staining.

For OCT4 staining, cells were PBS washed three times and fixed for 1 hour at 4° C with Paraformaldehyde, 16% w/v aq. soln., methanol free (ThermoFisher Scientific, 043368-9M) diluted to 4% in PBS. Cells were washed three times with PBS following fixation and blocking was performed in 2% BSA (Sigma-Aldrich, A2153-50G) with 0.1% Triton-X 100 (Millipore Sigma, 1.08603.1000) for 1 hour at room temperature. Primary antibody (goat anti-OCT-3/4, Santa Cruz sc-8628) was diluted 1:2500 in blocking buffer and 500µL was added to each well overnight at 4°C. Primary antibody was removed, and cells were washed three times with PBS. Secondary antibody (chicken anti-goat Alexa 647, Invitrogen A-21469) was diluted 1:1000 in blocking buffer and incubated overnight at 4°C. Cells were then washed three times with PBS and incubated with 300µL per well of Hoechst 33342 (Thermo Scientific, 62249) diluted 1:5000 in PBS for 5- 10 minutes at room temperature. Cells were washed once with PBS then stored at 4°C protected from light and evaporation until ready to image. OCT4 nuclear intensity was quantified using Fiji. Briefly, single slice confocal images were taken at the center of the sample using a 20x 0.7 NA objective (Nikon MRH48250). Once loaded into Fiji, individual color channels were split, and a threshold was imposed on the Hoechst channel using an Otsu range of 130-65535. These settings were used to convert the Hoechst channel to a Binary mask which was then adjusted with the ‘fill holes’ command and individual nuclei separated from each other with the ‘watershed’ command (both of which can be found in the Process > Binary dropdown menu). After visual inspection to assess accuracy of the mask, ‘Analyze Particles’ was run with size range set to 25-500 pixels (to remove small debris or ineffectively segmented nuclei from analysis) and 0-1 circularity. This mask was then applied to the OCT4 channel and the average measurement for each segmented nucleus was calculated to report across groups. Additional representative images were captured using an epifluorescence microscope (EVOS M5000 Imaging system); with 4x/10x phase ring, DAPI, Cy5, and 10X Objective, fluorite, LWD, phase-contrast, 0.30NA/7.13WD (Thermo Fisher, AMEP498).

For flow cytometry assessment, cell suspension was spun down at 300g for 4 minutes and resuspended in 1mL PBS for staining with 1 µL LIVE/DEAD^TM^ Fixable Orange (602) Viability Kit for 30 minutes at room temperature. The cells were pelleted, washed once with PBS, and then resuspended in 98 µL of PBS with 5% BSA (Millipore Sigma, A1595). The samples were then stained with 2 µL of Anti-human SSEA-4 APC-Vio® 770, REAfinity™ (Miltenyi Biotec) and incubated in the dark for 15 min at 4° C. Cell suspensions were then diluted in 1 mL of cold PBS and pelleted for 10 min at 400 g and 4° C, resuspended in PBS, filtered and analyzed by flow cytometry.

### Flow cytometry

Flow cytometry experiments were conducted on an Attune NxT Acoustic Focusing Cytometer using a 488 nm laser, 510/10 filter, and a voltage gain of 240 for eGFP fluorescence, a nm laser, 620/15 filter and a voltage gain of 400 for mOrange live/dead fluorescence, and a nm laser, 780/60 filter and a voltage gain of 400 for SSEA-4-APC-Vio7 fluorescence. FSC-A and SSC-A were used to gate cells, with FSC-H and FSC-A used to gate singlets. eGFP^+^, mOrange live/dead^-^, and SSEA-4-APC-Vio7^+^ gates were set using an untransfected and unstained negative control sample. Gating and analysis were conducted using FlowJo. The gating strategy and representative histogram plots are presented in Figure S1. Raw .fcs files will be made available on GitHub.

### hiPSC suspension spheroid culture

hiPSCs (37-1189) were seeded onto Matrigel® (Corning, 354277) in DMEM/F12 (Gibco, 11320033) coated 24-well plate transfected as described above and maintained in 1 µg/mL puromycin (Thermo Scientific, J67236.XF) in StemFlex Media (ThermoFisher, A3349401) for 5 to 7 days depending on growth rate and required cell number. To seed into suspension, cells were washed twice with D-PBS and incubated with Stempro^TM^ Accutase^TM^ (ThermoFisher Scientific, A1110501) for 3-5 minutes at 37°C. Dissociated cells were gently triturated to singularize and added to a StemScale media volume that is 4 times that of Accutase to neutralize dissociation. Cell suspension was then centrifuged at 200 g for 4 minutes to pellet cells. Supernatant was aspirated and cell pellet was resuspended in StemScale^TM^ PSC Suspension Medium (ThermoFisher Scientific, A4965001) media supplemented with Y-27632 (Millipore Sigma, 5092280001 or StemCell Technologies, 72304) to a final concentration of 10 µM. Live cells were counted using a Countess 3 (Invitrogen) cell counter with Trypan Blue Stain (Invitrogen, T10282) and added to a non-tissue culture treated 6 well plate at 300,000 cells per well in StemScale media with 10 µM Y-27632. Plates were then incubated on an orbital shaker (Thermo MaxQ 2000 CO2 Plus) rotating at 70 rpm. Half media changes with StemScale were performed the day after seeding and every other day following by tilting the plate at a 45° angle, allowing the spheroids to settle, and gently aspirating and replacing half the media. Puromycin selection was withdrawn for seeding and optionally added back the day following seeding using a complete media change also performed by tilting the plate to a 45° angle a pipetting off and replacing the entire media volume.

### saRNA-based differentiation of hiPSCs to neurons

Transfection of hiPSCs (iPS11) plated on Laminin (StemCell Technologies, 200-0117) coated 24-well plates with 500 ng of Ngn2-2A-eGFP saRNA was conducted as described above, except using a cell density of 200,000 cells/24-well (∼60% confluency) and the 4 hour media refresh was performed using neural induction media (DMEM/F12 supplemented with 1:100 Glutamax™,1:100 Non-essential amino acids (Gibco), 1:100 N2 supplement (Gibco), and 1:100 Penicillin/streptomycin(Gibco)). One day post-transfection, media was manually aspirated gently with a P200 pipette and replaced with 500 µL of inducttion media containing 1 µg/uL Puromycin. Two days post-transfection, media was manually aspirated with a P200 pipette, cells were gently washed with PBS, and the well refreshed with neural maturation media (DMEM/F12 supplemented with Glutamax™ supplement (100x, Gibco), 1:100 Non-Essential Amino Acids (Gibco), 1:100 N2 supplement (Gibco), 1:50 B27 supplement (Gibco), BDNF (10 ng/ml, R&D Systems), NT3 (10 ng/ml, Peprotech), and 1:100 Penicillin/Streptomycin) containing 1 µg/uL Puromycin. Starting three days post transfection, daily half media changes were made with 250 µL maturation media without Puromycin.

### Fixation, immunofluorescent staining, and purity quantification of induced neurons

For induced neurons, immunostaining was carried out at 24-well scale at 6 dpt of Ngn2-2A-eGFP. All wash/addition steps were carried out via manual aspiration (i.e. slowly removing or adding media with a P200 pipette), rotating the quadrant the pipette tip was submerged clockwise for each pipetting step to avoid over pipetting in one area of the well. Culture media was removed from iNs and cells were washed three times with PBS. Cells were fixed with 4% paraformaldehyde for 1 hour at 4°C. After 1 hour, the fixative was removed, and cells were washed three times with cold PBS. Cells were then blocked and permeabilized with 5% FBS and 0.5% Tween-20 in PBS for 24 hours at 4°C. Following permeabilization and blocking, primary antibodies, either mouse anti-TUJ1 (BioLegend 801202) or rabbit anti-MAP2 (Cell Signaling Technologies 4542) were added at a 1:500 dilution in blocking buffer (5% FBS, 0.1% Tween-20 in PBS) and incubated overnight at 4°C. Cells were washed three times with 0.1% Tween-20 in PBS and blocked and permeabilized in 5% FBS and 0.5% Tween-20 in PBS for 1 hour at 4°C. DAPI (1x) (Tocris 5748) and secondary antibodies, either goat anti-mouse Alexa Fluor 555 (Invitrogen, A-21422) or donkey anti-rabbit Alexa Fluor 647 (Jackson ImmunoResearch Laboratories, 711-605-152) were added at a 1:500 dilution in blocking buffer and incubated at room temperature for 1 hour. Cells were washed with 0.1% Tween-20 in PBS three times and cells were kept in PBS following the last wash. Images and visualization were carried out with the Keyence™ All-in-one fluorescence microscope BZ-X800.

To quantify the purity of the induced neurons, five sections of a DAPI and TUJ1-stained well were randomly sampled and imaged. DAPI stained nuclei were segmented with CellProfiler [77], using the IdentifyPrimaryObjects module. This module set the object diameter to a Min of 8 pixels and a Max of 50 pixels, implemented an adaptive Minimum Cross-Entropy thresholding method with a smoothing scale and correction factor of 1.65 and 1, respectively, and used pixel intensity to distinguish clumped objects. The pipeline is provided in a .zip folder on GitHub. After segmentation to obtain the total number of nuclei in the image section, the number of TUJ1-negative nuclei (corresponding to residual stem cells) were manually counted and subtracted from the total nuclei count to determine the number of TUJ1-positive nuclei (corresponding to the number of iNs). TUJ1-positive nuclei and total nuclei across all five-image section were summed and purity was calculated using the ratio of TUJ1-positive nuclei to total nuclei. An example of the segmentation strategy for one technical replicate is provided in Figure S5.

### Quantitative PCR analysis of induced neurons

Three separate differentiation experiments were conducted using iPS11. For each replicate, induced neurons obtained from three 24-wells transfected with Ngn2-2A-eGFP saRNA and negative control cells obtained from one 24-well transfected with eGFP saRNA were harvested six days post transfection, resuspended in 200 µL of RNA later buffer, and stored at – 80° C. RNA from all three experiments were isolated simultaneously using the Monarch Total RNA Miniprep Kit (New England Biolabs, T2010) with an additional on-column Dnase I (New England Biolabs, M0570) treatment step. RNA samples were eluted with 50 µL of nuclease-free water. cDNA was synthesized from 6 µL of eluted RNA using the ProtoScript First Strand cDNA Synthesis Kit (New England Biolabs, E6300) using oligo-dT primers. cDNA samples were stored at −20°C until qPCR.

qPCR was performed at the MIT BioMicro Center on a Roche LightCycler 480 with three technical replicates per condition. Reaction mixes were assembled using 5.0 µL KAPA SYBR FAST qPCR Master Mix (2X) Universal (Kapa Biosystems, KK4600), 0.5 µL 2 µM forward and reverse primers, 0.5 µL cDNA product, and 1.5 µL nuclease-free water. The primer sequences used for each gene target are provided in the supplementary information. Using the “High Sensitivity” analysis mode, Ct values were called for each technical replicate, with average Ct values determined by calculating the arithmetic mean of the three technical replicates. 2^-ΔCt^ values were calculated relative to the RPL37A levels for each sample and log10 transformed.

### Imaging of HyPer7 reporter

Generation of the HyPer7-Mito saRNA and transfection into hiPSCs was performed as described above. Two days post transfection, MitoTracker™ Deep Red FM dye (Thermo Fisher Scientific, M22426) was added to the cells at a concentration of 50nM and incubated at 37°C for 15 minutes. Culture medium was replaced with 1.2 mL of HBSS solution (Thermo Fisher Scientific, 14025092) supplemented with 20 mM HEPES (Thermo Fisher Scientific, 15630080). D-alanine (TCI, A0177) or Hydrogen Peroxide (H202) (Thermo Fisher Scientific, H325-500) were added to the appropriate conditions to a final concentration of 40mM and 25μM, respectively. Imaging was performed using an Inverted Evident FluoView FV4000 microscope with a 60x/1.30 silicon immersion UPlan Superapochromat objective. The HyPer7 fluorophore was excited sequentially with 405 nm and 488 nm lasers, and emission was detected with a 520nm filter. The HyPer7 ratio was calculated using the Fiji distribution of ImageJ by performing background subtraction, then dividing the mean intensity of each 488nm excitation image (oxidized HyPer7) by the corresponding 405nm excitation image (reduced HyPer7). Colorized ratio images were generated as previously described [78], [79]. Briefly, each 488nm excitation image was divided by its corresponding 405nm excitation image, and the resulting image was depicted in the “Fire” lookup table.

### Voltage and calcium imaging and analysis in 2D cell culture

Mouse hippocampal cell culture was prepared at 24 well scale as described above and were transfected with 500 ng of saRNA diluted in Opti-MEM™ (ThermoFisher Scientific, 31985070) to a volume of 25 µL. In parallel, 1 µL of Lipofectamine™ MessengerMAX Transfection Reagent (ThermoFisher Scientific, LMRNA008) was added to 24 µL Opti-MEM™. The Lipofectamine™ mixture was preincubated for 15 minutes before adding the diluted saRNA, and the saRNA/ Lipofectamine™ mixture was incubated for an additional 5 minutes. After the final incubation step, the saRNA/LNP complexes were added drop wise to the well in 500 µL of maintenance media, and the plate was gently rocked and returned to the incubator. After 4 hr, 500 µL of additional maintenance media was added to each transfected well. hiPSC-derived CMs were prepared at 96 well scale and were transfected with 100ng saRNA in 5 µL Opti-MEM^TM^ complexed with 0.2 µL Lipofectamine™ MessengerMAX Transfection Reagent in 4.8 µL Opti-MEM^TM^ in 100 µL maintenance media. Refreshed with an additional 100µL after 4 hours. A complete media change was performed on transfected wells the following day. Neural cultures were transfected at ≥ 14 days *in vitro* and hiPSC-derived cardiomyocytes were transfected once beating resumed following replating at day 14 (between 4 and 5 days).

Live neuron and cardiomyocyte imaging was performed in their maintenance media at room temperature on a spinning disk confocal microscope (a Yokogawa CSU-W1 Confocal Scanner Unit on a Nikon Eclipse Ti microscope) equipped with a 40X 1.15 NA water immersion objective (Nikon MRD77410) and a Zyla PLUS 4.2 Megapixel camera controlled by NIS-Elements AR software. The filter set for GFP (488nm excitation; FITC emission filter) was used for imaging ASAP5 and the filter set for mRuby (561nm excitation; Cy3 emission) was used for imaging jRCaMP1b. Voltage imaging (ASAP5) was performed at 300 ms exposure (3.33 Hz) for neuronal cell types and 100 ms exposure (10 Hz) for cardiac cell types. Calcium imaging (jRCaMP1b) was performed at 100 ms exposure (10 Hz) for neuronal cell types and 100 ms exposure (10 Hz) for cardiac cell types. For each sample, imaging was performed on a single focal plane near the mid-section of cells. hiPSC-derived CMs with beating patterns that kept the cell body in focus were selected for representative traces. From primary mouse hippocampus cultures, cells with characteristic neuronal morphology, including small cell bodies and long, branched processes, were visually identified and treated as putative neurons. Sliding window normalization was applied to correct for slow baseline signal drift. Briefly, for each recorded trace, the raw fluorescence intensity vector was extracted, and a baseline fluorescence (F_!_) was estimated using a sliding window set between 15 and 100. Within each window, the baseline was defined between the 10^th^ and 50^th^ percentile (median) of fluorescence values, providing an estimate against noise and transient fluctuations. The baseline trace was then interpolated across the full recording interval to generate a continuous estimate of F_!_ (defined as the value under which P percent of fluorescence values fall in W frame period). The top percentile values (50^th^ to 90^th^) were used to calculate baseline of ASAP5 traces (fluorescence decreases in response to a voltage change) and inverted for all visual displays. Each sensor was imaged 2 days post transfection (with a media change at 24 hours) and representative traces for three cells are shown in Figure 3.

Normalized fluorescence was calculated as (F(t) − F_!_(t))/F_!_(t), where F(t) is the raw trace and F_!_(t) is baseline calculated with sliding window normalization at each time (t). Both raw and baseline traces were plotted for visual inspection, and normalized traces were displayed to quantify relative fluorescence changes (noting that the polarity is inverted for inverse indicators such as ASAP5). The normalized traces, raw traces, and calculated baseline fluorescence with accompanying window size and percentile settings are presented in Figure S7.

### Differentiation of hiPSCs to cardiomyocytes in suspension

hiPSC spheroids were differentiated according to manufacturer instructions for cardiomyocyte differentiation from PSC suspension culture in StemScale^TM^ PSC Suspension Medium (ThermoFisher Scientific, A4965001) [80]. Briefly, upon reaching an average diameter of between 250 µm and 300 µm (between 4 to 5 days), as assessed by brightfield or phase contrast microscopy, wells were washed with RPMI 1640 medium (ThermoFisher Scientific, 11875093) and incubated for 48 hours in RPMI supplemented with B-27^TM^ Supplement, minus insulin (ThermoFisher Scientific, A1895601) and 6 µM CHIR-99021 (Selleck Chemicals, S2924) (day 0). Exactly 48 hours following CHIR-99021 addition (day 2), wells were washed with RPMI and incubated for 48 hours in RPMI with B-27 minus insulin and 7.5 µM IWP-2 (Fisher Scientific, 35-331-0). Following IWP incubation (day 4), all wells were changed into RPMI with B-27 minus insulin for 48 hours. On day 6, wells were changed into RPMI media supplemented with B-27^TM^ Supplement (50X), serum free (ThermoFisher Scientific, 17504044). 50% media changes in RPMI with B-27 were performed every other day following day 6. All incubations were conducted at 37°C with 5% CO2 on a plate shaker rotating at 70rpm.

### Fixation and immunofluorescent staining of cardiac spheroids

For cardiac spheroids, immunostaining was carried out at 24-well scale on day 14 and day 30 following differentiation induction at day 0. Cardiac spheroids were collected in an Eppendorf tube and kept on ice until fixation. Spheroids were allowed to settle by gravity sedimentation (∼1 minute) and were washed with 1 mL PBS prior to fixation for 30 minutes with Paraformaldehyde (PFA), 16% w/v aq. soln., methanol free (ThermoFisher Scientific, 043368-9M) diluted to 4% in PBS with gently agitation. Following incubation, PFA was removed, and cardiac spheroids were washed for 5 minutes in PBS three times. Cardiac spheroids were then transferred to a 1% BSA (Millipore Sigma, A1595) coated 24 well plate and blocked with 1% BSA and 0.5% Triton X-100 (Merck, 108603 1000) in PBS for 2 hours at room temperature or overnight at 4°C. Primary antibodies (mouse anti-cTnT – Abcam, ab8395; rabbit anti-nkx2.5 – Cell Signaling Technology, #8792) were diluted in 1% BSA in PBS according to manufacturer recommendations (1:1000) and added for 4°C with gently agitation for 2 days. Secondary antibodies (goat anti-mouse 594 – Invitrogen, A11005; chicken anti-rabbit 647 – Invitrogen, A21443) were diluted to 1:200 and added after three 10-minute washes with PBS. Following 2-day incubation at 4°C with gently agitation, samples were washed three times with PBS and incubated overnight at 4°C with Hoechst (Thermo Scientific, #62249) diluted to 1:5000 in PBS. Samples were then washed and stored in PBS.

### Quantification of labeling percentage in cardiac spheroids

Cardiac spheroids were fixed and stained as described above. 4-channel z-stacks were taken at a step size of 12.5 µm through the depth of spheroids using a spinning disk confocal (a Yokogawa CSU-W1 Confocal Scanner Unit on a Nikon Eclipse Ti microscope) equipped with a 4X 0.2NA objective (Nikon MRD70040) and a Zyla PLUS 4.2 Megapixel camera controlled by NIS-Elements AR software. Hoechst: 405 nm excitation, 500 ms exposure time, 70.8 laser intensity; eGFP– 488 nm excitation, 300 ms exposure time, 100.0 laser intensity; NKx2.5 – 640 nm excitation, 100 ms exposure time, 100.0 laser intensity; cTnT – 561 nm excitation, 600 ms exposure time, 100.0 laser intensity.

Maximum intensity projections were generated for each image and individual spheroids were segmented based on the Hoechst channel in Fiji. Briefly, images were split into individual color channels and maximum projections were generated. A manual threshold was set on the Hoechst channel to segment spheroids from background and was converted to a binary mask and ‘analyze particles’ was used to generate a set of individual ROIs that represent each spheroid. Spheroids that were negative for cTnT signal were manually excluded from analysis as not having undergone differentiation. The eGFP maximum projection was converted to 8 bit and a histogram of pixel values were generated for each ROI with 256 bins. The percentage of pixels that are in bins 26 through 256 of each histogram out of the total number of pixels is used to represent the % each spheroid that is eGFP positive.

### Calcium imaging in cardiac spheroids

Calcium imaging in cardiac spheroids was performed at room temperature in RPMI 1640 medium (ThermoFisher Scientific, 11875093) B-27^TM^ Supplement (50X), serum free (ThermoFisher Scientific, 17504044). Using a 4x 0.2 NA (Nikon MRD70040), 10x 0.45 NA (Nikon MRD70170), or 20x 0.7 NA (Nikon MRH48250) objective using 561nm excitation and Cy3 emission filter settings and 300 ms exposure time (3.33Hz).

50 Hz imaging was performed at 37°C on an epifluorescence microscope (a Lumencor SPECTRA X Light Engine on a Nikon Eclipse Ti-2 microscope) with a 20x 0.80 NA objective (Nikon MRD70270) and ORCA-Fusion BT Digital CMOS camera (Hamamatsu C15440-20UP) controlled by NIS-Elements software. Images were captured with 20 ms exposure time using Texas Red excitation and filter settings.

### Calculation of calcium peak parameters in cardiac spheroids

In Fiji, an ROI was manually drawn to encompass regions of the image that showed changes in fluorescence and values were measured throughout the z-stack (representing time). Each fluorescence time series representing calcium dynamics were analyzed using a custom MATLAB function (made available on GitHub). Briefly, raw traces consist of a time vector (t) and normalized fluorescence intensity values (y, scaled between 0 and 1 to account for differing baseline brightness to directly compare only temporal parameters). For acquisition rates lower than 30 fps (most taken at 3.33 Hz unless otherwise specified), traces were upsampled by a piecewise cubic Hermite interpolation (pchip) to achieve an effective sampling frequency of > 30 Hz. Peaks were identified using the findpeaks function with a minimum peak prominence threshold of 0.2 (out of 0 to 1) and a minimum inter-peak interval of 0.3 seconds. These plots were then displayed graphically to ensure accurate peak detection by visual inspection. For each 30 second trace, the total number of detected peaks was recorded. Inter-beat intervals (IBIs) were calculated as the time differences between successive peaks, and instantaneous beat frequencies were expressed as beats per minute (BPM = 60/IBI). Beat-to-beat variability was quantified as the coefficient of variation of the IBI (CV_IBI). Calcium traces with 2 or fewer identified peaks were removed from analysis since they falsely yield a coefficient of variation of 0. Because the original sampling interval imposes a quantization error (0.3/√12 ≈ 0.087 sec based on the standard deviation of a uniform distribution), we estimated residual timing uncertainty after interpolation (0.02–0.03 s depending on SNR) which propagates to the estimated BPM uncertainty. For example, if peaks are detected 3 seconds apart, ±0.03 second results in ±1% uncertainty in BPM. Only for 50 Hz sampled traces were rise kinetics calculated (>10 frames per risetime without up-sampling) while decay kinetics (which last approximately 1 second) were calculated for all traces (points along observation curve are shown in Figure S8). For the largest detected peak, the baseline was defined as the minimum fluorescence value preceding the peak. The 10%, 50%, and 90% fluorescence values relative to the peak amplitude were then computed. Rise time was measured as the time interval between the first crossing of the 10% and 90% fluorescence levels prior to the peak maximum. Decay was quantified as the time required for the signal to decrease from peak amplitude to 50% of the peak amplitude (T50) following the peak maximum. All extracted metrics were compiled for each trace.

### Statistical analysis

Each individual data point on summary graphs represents a biological or technical replicate and are specified as such in figure legends. Unless specified otherwise, data are reported as arithmetic means ± standard error of the mean (SEM) when reporting variation between averaged values and arithmetic means ± standard deviation when whole distributions are reported. Presented statistical analyses of normally distributed data include Welch’s Student T test for pairwise comparisons and Ordinary one-way ANOVA with Bonferroni correction for multiple comparisons. Statistical comparisons of non-normally distributed data with two groups represent a Mann-Whitney U test. All calculations were performed using GraphPad Prism v.8 or MATLab version 2023a, with significance set at ns: p ≥ 0.05, *: p < 0.05; **: p < 0.01; ***: p < 0.001; ****: p < 0.0001.

