## Supplemental Information for "Self-amplifying RNA enables rapid, durable, integration-free programming of hiPSCs"

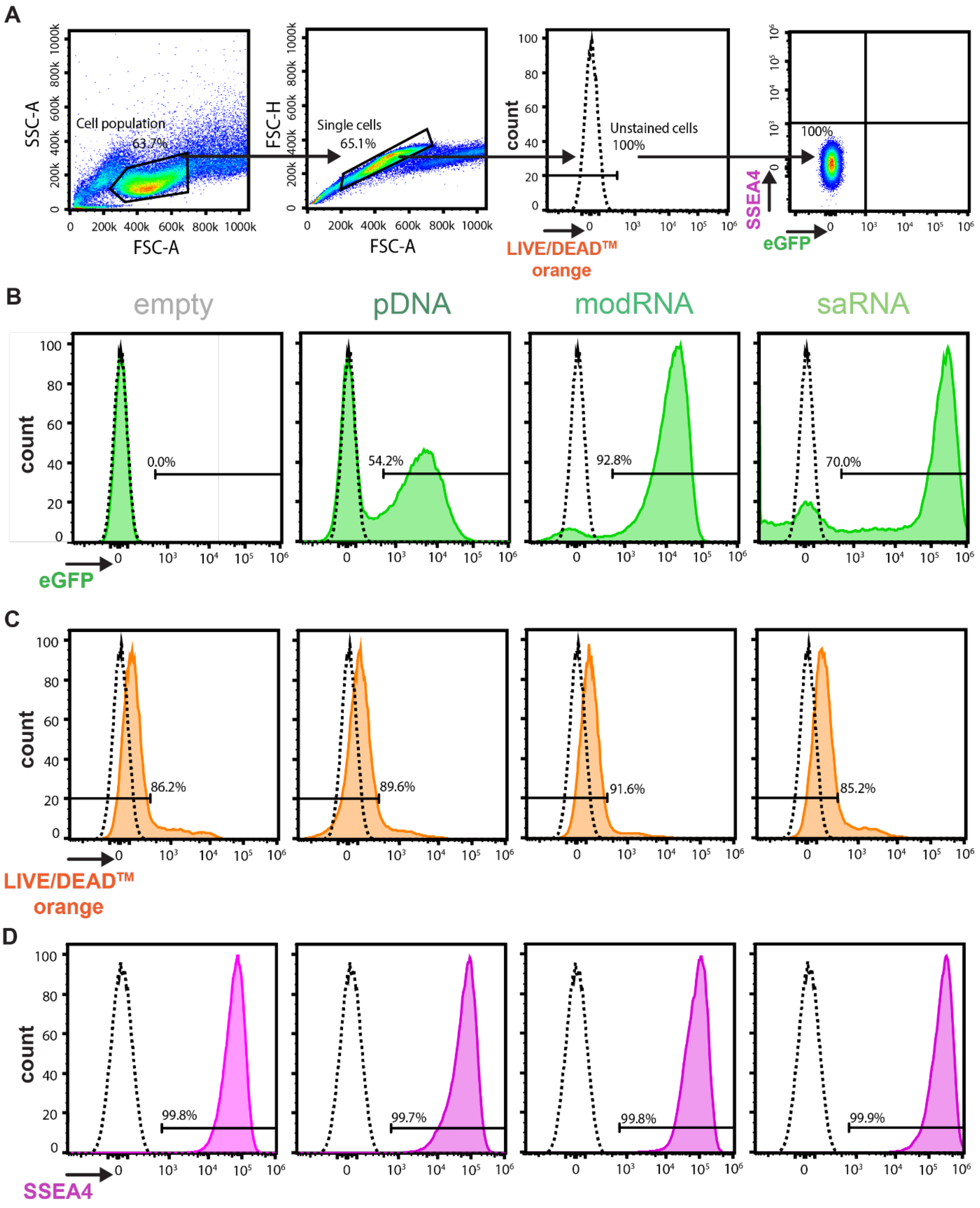


**Figure S1. saRNA enables sustained transgene expression in 2D and 3D hiPSC cultures. Related to Figure 1 and Methods. A.** Dot plots demonstrating flow cytometry analysis gating strategy. Forward and side scatter were used to identify the cell population, with forward scatter area versus height used to isolate the singlet population. An unstained, untransfected condition (dashed line) was used to set gates for the mOrange live/dead stain negative population, which was then used to set gates for the Anti-SSEA-4-APC-Vio770 negative and eGFP negative populations. **B**-**D.** Representative histograms for empty, pDNA, modRNA, and saRNA transfection conditions for **B.** eGFP **C.** mOrange live/dead and **D.** Anti-SSEA-4-APC-Vio770.

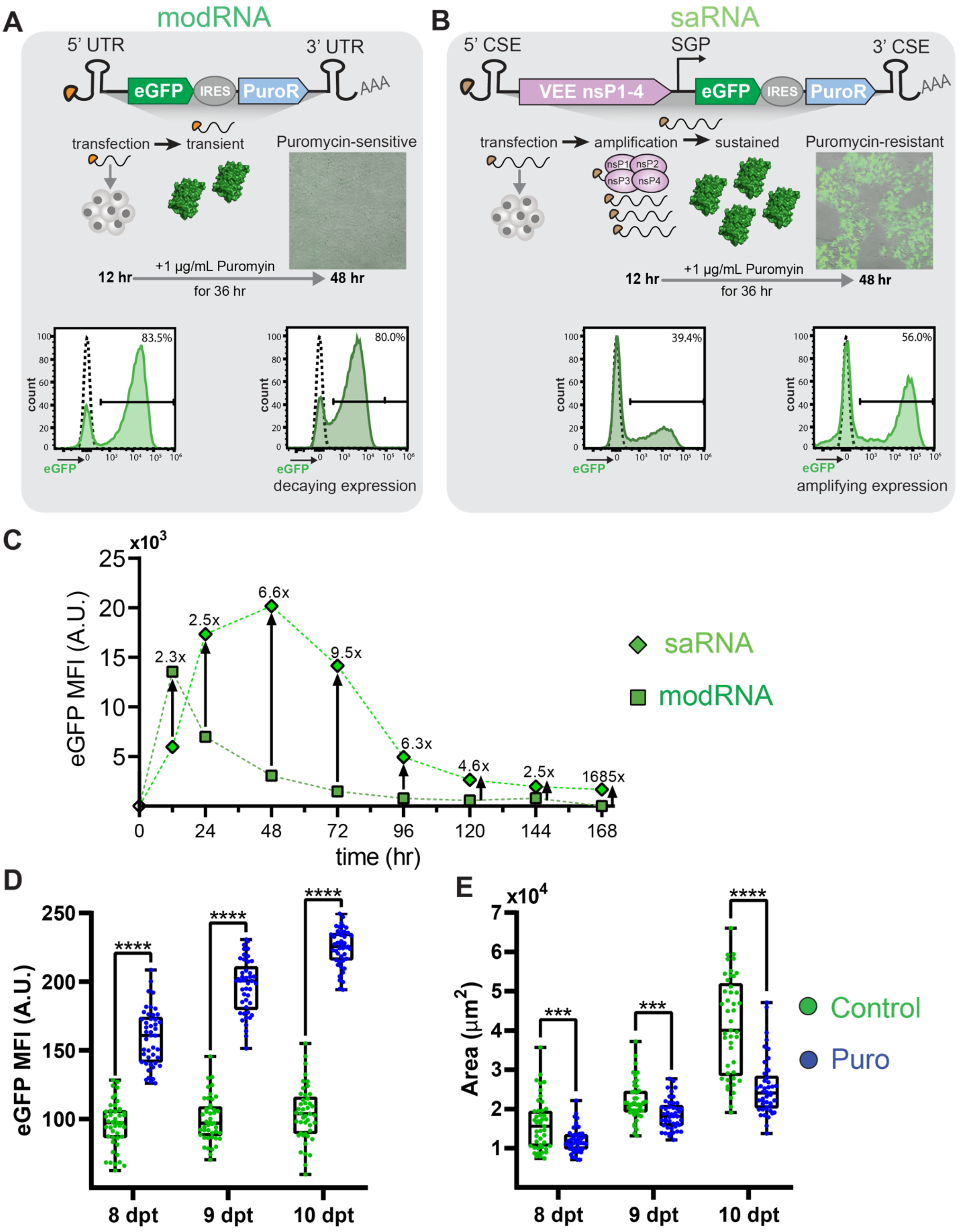


**Figure S2 saRNA enables sustained transgene expression in 2D and 3D hiPSC cultures. Related to Figure 1 and Methods.** The sustained expression from saRNA compared to modRNA potentiates nonintegrating, longer-term maintenance of transgenic cargoes. **A.** Delivery of modRNA encoding an eGFP-IRES-PuroR transgenic cassette fails to confer puromycin resistance when administered for 36 hr at 12 hr post-transfection, attributable to transgene expression decay at 48 hr compared to 12 hr (measured via flow cytometry). As the puromycin-treated condition resulted in complete cell death, the 48 hr flow cytometry histogram is representative of modRNA-transfected cells not treated with puromycin. **B.** Delivery of saRNA encoding the same eGFP-IRES-PuroR transgenic cassette robustly confers puromycin resistance, attributable to transgene expression amplification at 48 hr compared to 12 hr, (measured via flow cytometry). The cognate 48 hr flow cytometry histogram is representative of saRNA-transfected cells treated with puromycin for 36 hr at 12 hr post-transfection. **C.** An eGFP expression time-course comparison between modRNA and saRNA demonstrates the differential kinetics of modRNA (fast, transient expression) versus saRNA (relatively slower onset, more sustained expression). Cells transfected with either modRNA or saRNA were analyzed via flow cytometry at 12 hr, 24 hr, and every 24 hr thereafter until seven days post-transfection (168 hr). Cells analyzed at later timepoints were passaged at 72 hr and 144 hr post-transfection, consistent with a standard 3-day passaging schedule to prevent overcrowding. **D.** Average eGFP fluorescence and **E.** cross-sectional area for individual spheroids reported in Figure 1H. N = 45 per distribution (5 spheroids per image and 3 images per well for 3 total wells).

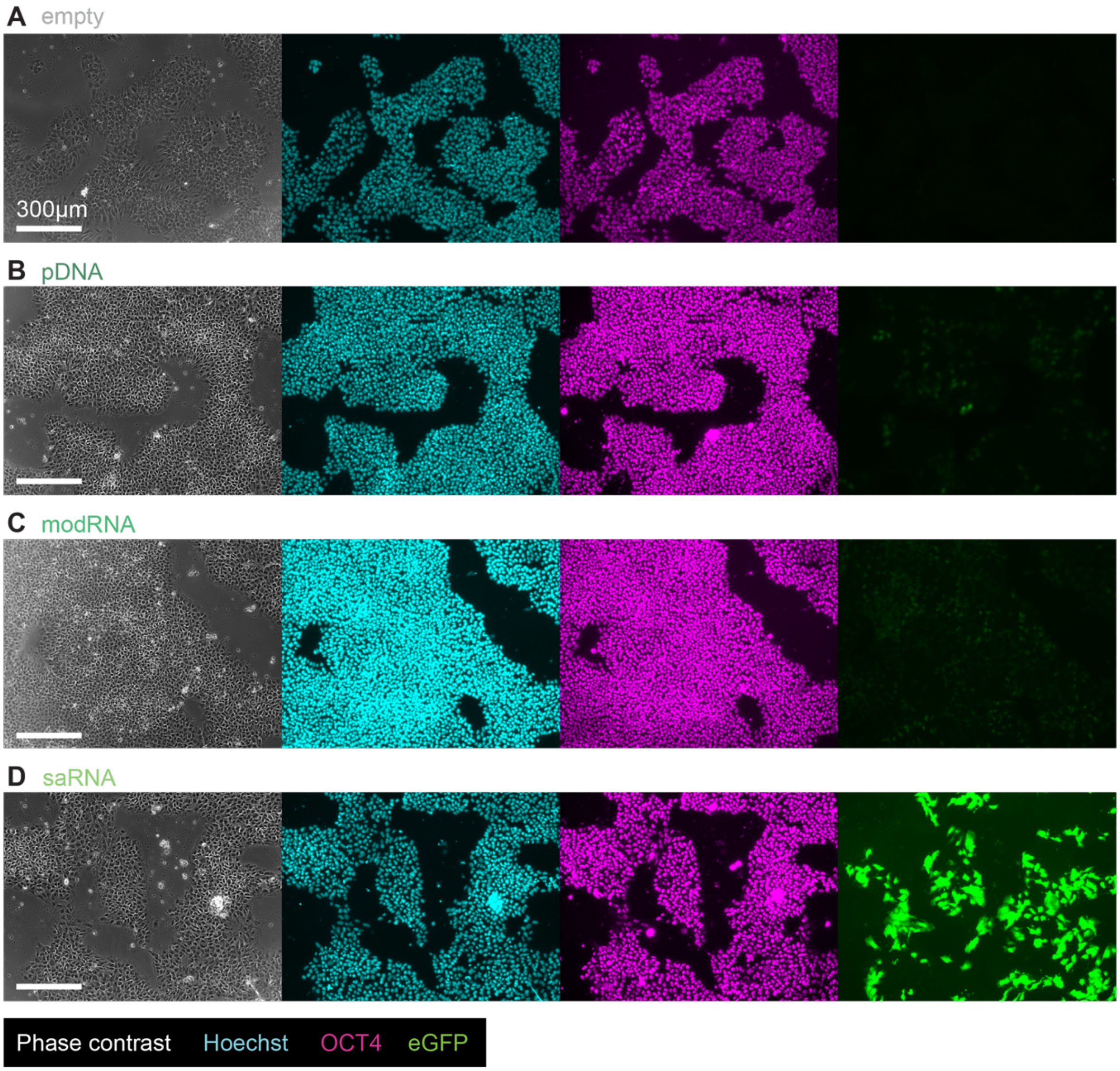


**Figure S3 saRNA does not impact cell morphology. Related to Figure 1 and Methods.** Representative epifluorescence images of hiPSC (iPS11) morphology in phase contrast, nuclei stained in Hoechst, OCT4 expression, and eGFP signal intensity for **A**. untransfected control hiPSCs and hiPSCs transfected with **B**. plasmid DNA, **C**. modified RNA, or **D**. self-amplifying RNA encoding the same eGFP-IRES-PuroR gene cassette. Cells were fixed 48 hours post transfection for staining.

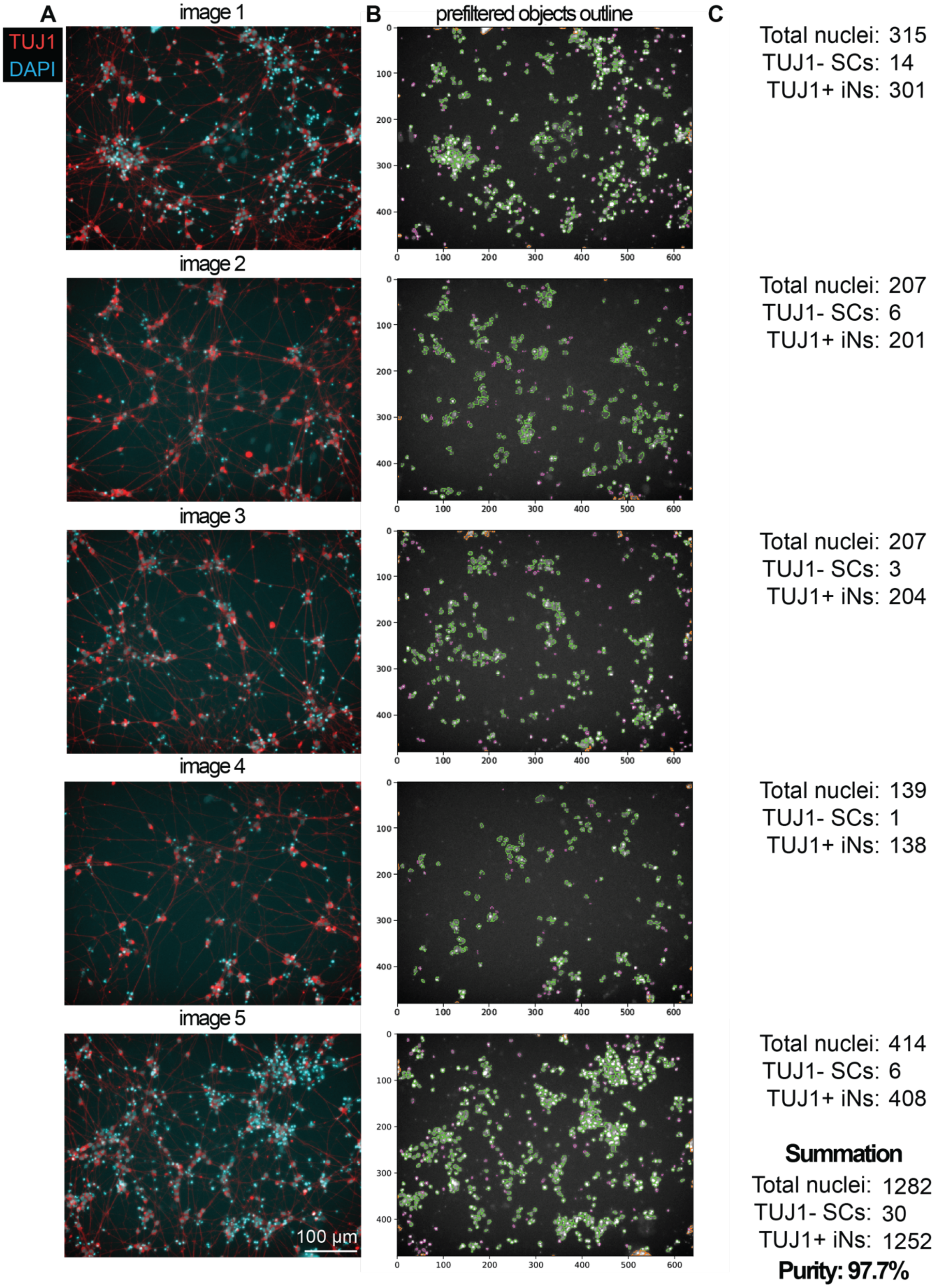


**Figure S4. saRNA-based delivery of Ngn2 efficiently differentiates hiPSCs to neurons. Related to Figure 2 and Methods. A.** Fluorescence microscopy images of random well positions used for iN purity quantification**.** Blue channel is DAPI and Red channel is TUJ1 **B.** CellProfiler segmentations of DAPI images shown in **A.** Green objects correspond to nuclei, and purple objects correspond to rejected objects. **C.** Total nuclei counts (determined by CelllProfiler), TUJ1- SCs (determined by manual counting), and TUJ1+ iNs (calculated by subtracting SC nuclei from total nuclei) for each section, with totals summed and used to calculate purity percentages.

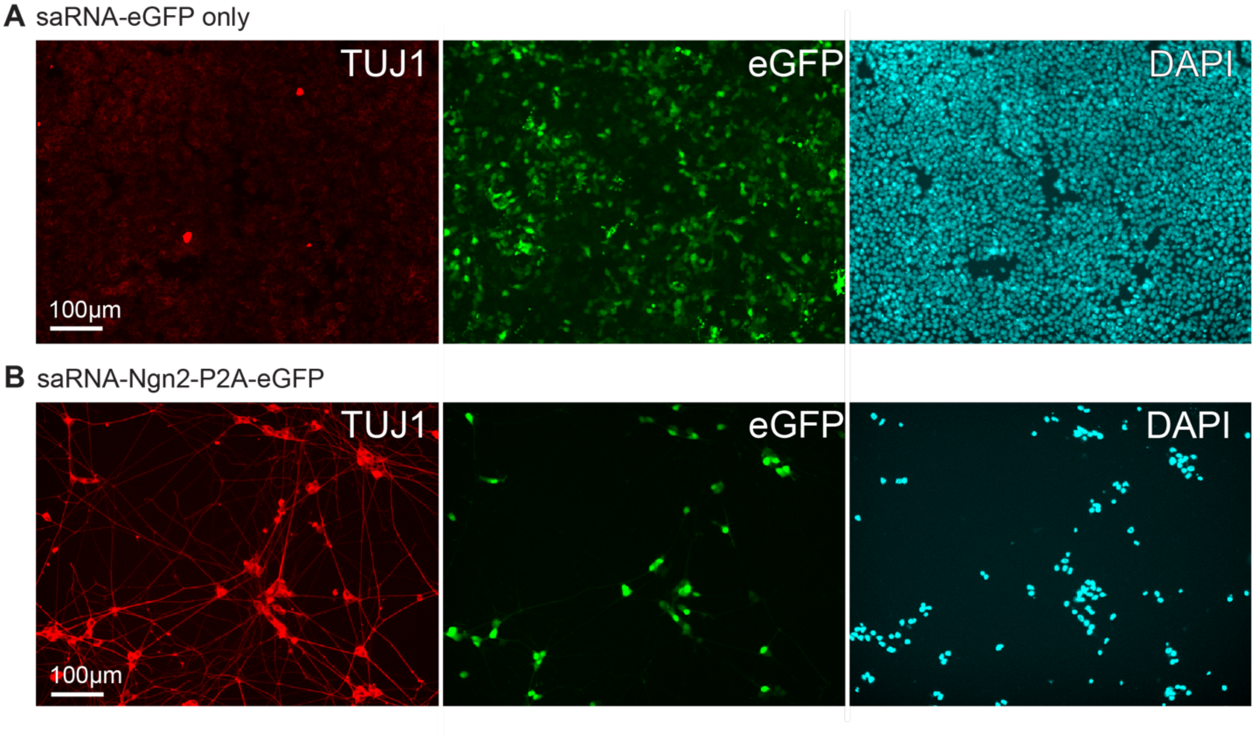


**Figure S5. Comparison of eGFP only versus Ngn2-P2A-eGFP saRNA in generating TUJ1 positive neuronal cells.** Representative image of cells fixed at day 6 with nuclei stained with DAPI, TUJ1 expression, and eGFP signal in **A.** hiPSCs transfected with eGFP carrying saRNA and subjected to neural induction and neural maturation medias in parallel to **B**. hiPSCs transfected with Ngn2-P2A-eGFP.

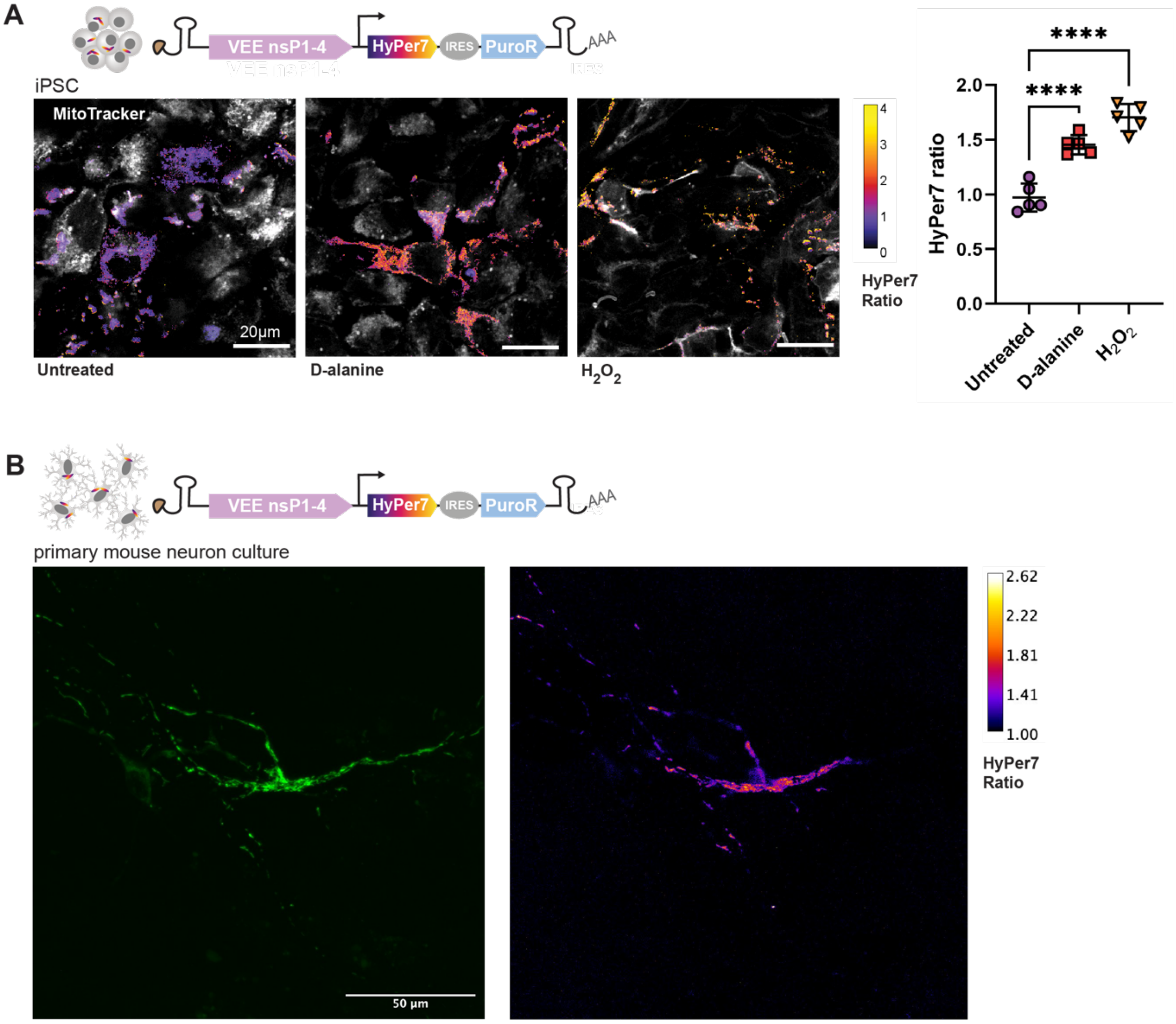


**Figure S6. saRNA-based delivery of HyPer7 to hiPSCs and neuronal culture. Related to Figure 3 and Methods. A.** HyPer7 reporter with N-terminal MLS (mitochondrial targeting signal of cytochrome c oxidase subunit VIII) in hiPSCs. The reporter contains D-amino oxidase, an enzyme which produces $H_2O_2$ upon addition of D-alanine. Ratio of oxidized (excitation at 488 nm) to reduced (excitation at 405 nm) state of Hyper7 is color-coded according to the accompanying color bar, where values closer to 4 indicate more $H_2O_2$ and closer to 0 indicate less. Mitotracker fluorescence is shown in gray. Scale bar 20 µm. Quantification of average HyPer7 ratio from 5 fields of view for untreated, 40 mM D-alanine, and 25 µM $H_2O_2$. **B.** HyPer7 expressed in primary mouse hippocampal cell culture 48 hours post-transfection. (Left) oxidized HyPer7 shown in green and (right) HyPer7 ratio imaged and calculated as described in A. Scale bar 50 µm.

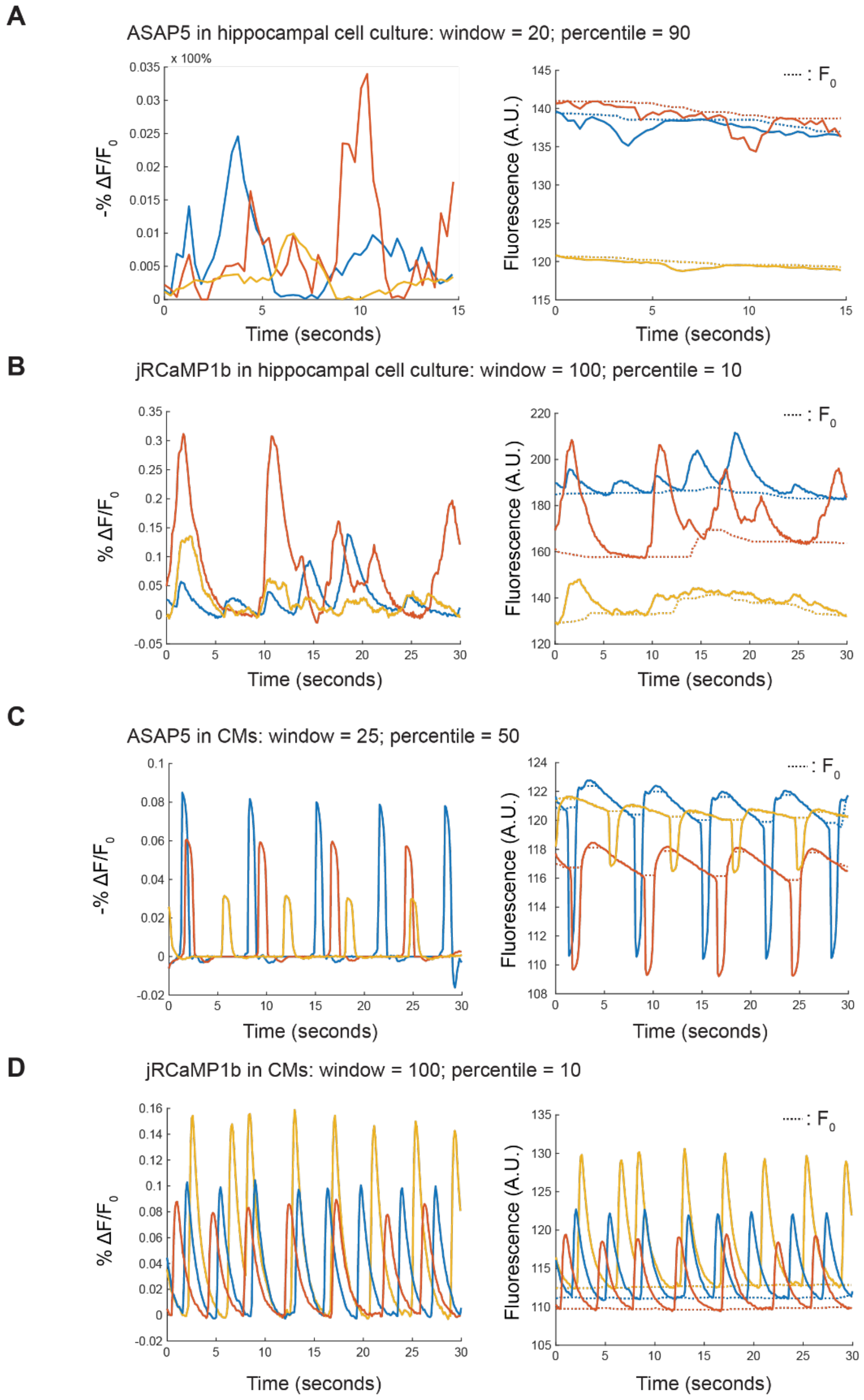


**Figure S7. Sliding window normalization of voltage and calcium traces in primary mouse hippocampus culture and hiPSC-derived cardiomyocytes. Related to Figure 2 and Methods.** 3 normalized ($|F-F_0|/F_0$) representative traces (blue, yellow, orange) per cell type and reporter combination where the accompanying plot on the right shows the raw fluorescence and the dotted line represents $F_0$ as calculated by sliding window normalization with window size (number of datapoints) and percentile values reported. **A.** ASAP5 voltage reporter in primary cell culture from mouse hippocampus with window size of 20 and percentile 90. **B.** jRCaMP1b calcium reporter in primary cell culture from mouse hippocampus with window size of 100 and percentile 10. **C.** ASAP5 voltage reporter in hiPSC-derived cardiomyocytes (CMs) with window size of 25 and percentile 50. **D.** jRCaMP1b calcium reporter in CMs with window size of 100 and percentile 10.

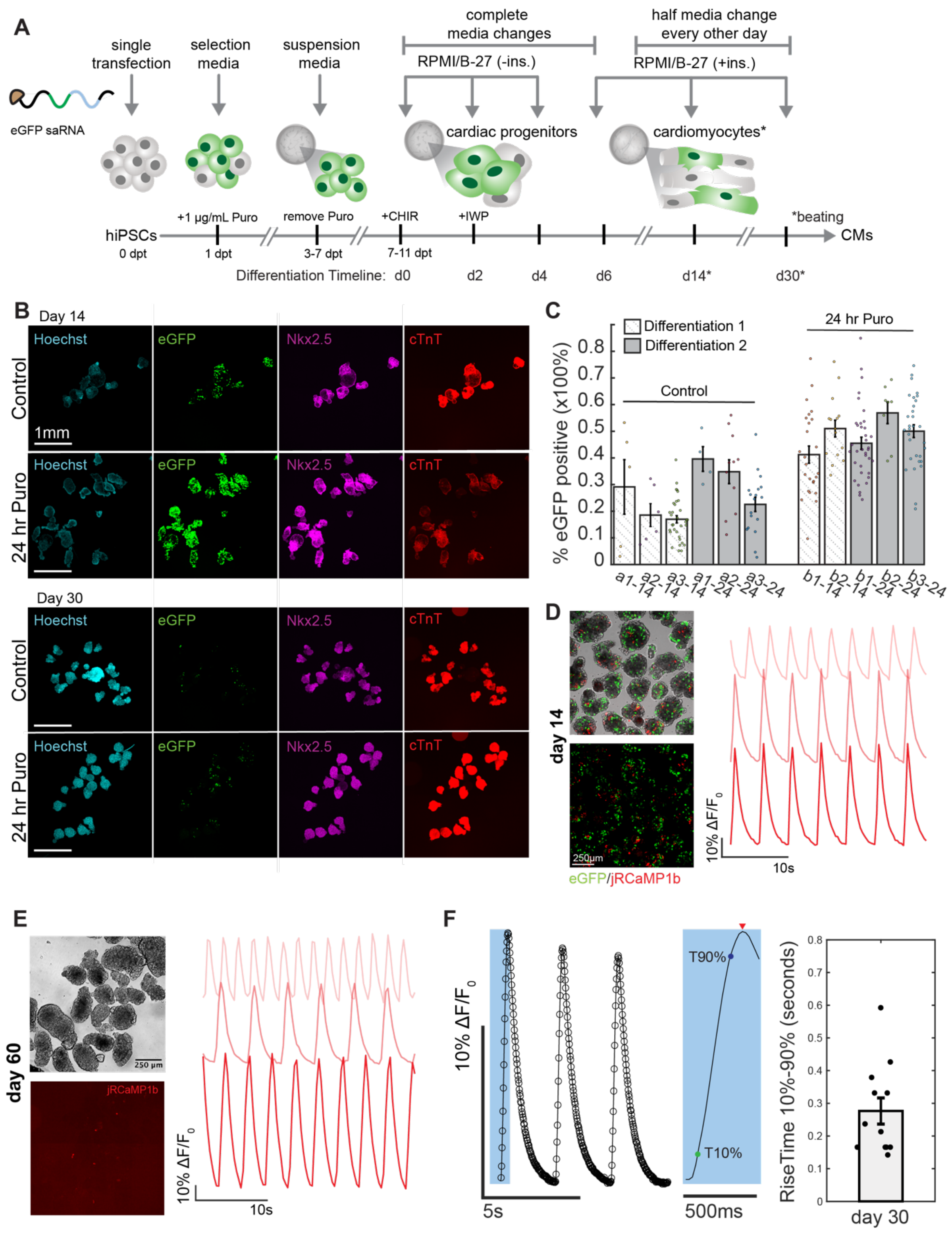


**Figure S8. saRNA-based delivery of fluorescent reporters to observe cardiac differentiation. Related to Figure 4 and Methods. A.** Detailed schematic of cardiac differentiation with saRNA constructs. **B.**

Representative IF images of day 14 and day 30 cardiac spheroids (with and without optional 24-hour puromycin selection following seeding into suspension) expressing saRNA-eGFP and stained for nuclear marker Nkx2.5 and sarcomere component cTnT. Related to Fig. 4B. **C.** Quantification across individual wells and differentiations related to Fig. 4C. **D.** eGFP and jRCaMP1b expressing cardiac spheroids and representative calcium traces at day 14 of differentiation. **E.** Representative brightfield and jRCaMP1b images with accompanying example calcium traces captured at day 60 of differentiation. **F.** Representative trace of day 30 differentiation captured at 50 fps imaging rate showing sufficient brightness to capture dynamics at lower exposure time. Inset showing rise time calculation for highlighted peak from time of 10% brightness to 90% brightness. Rise time is calculated for N = 11 cardiac spheroids.

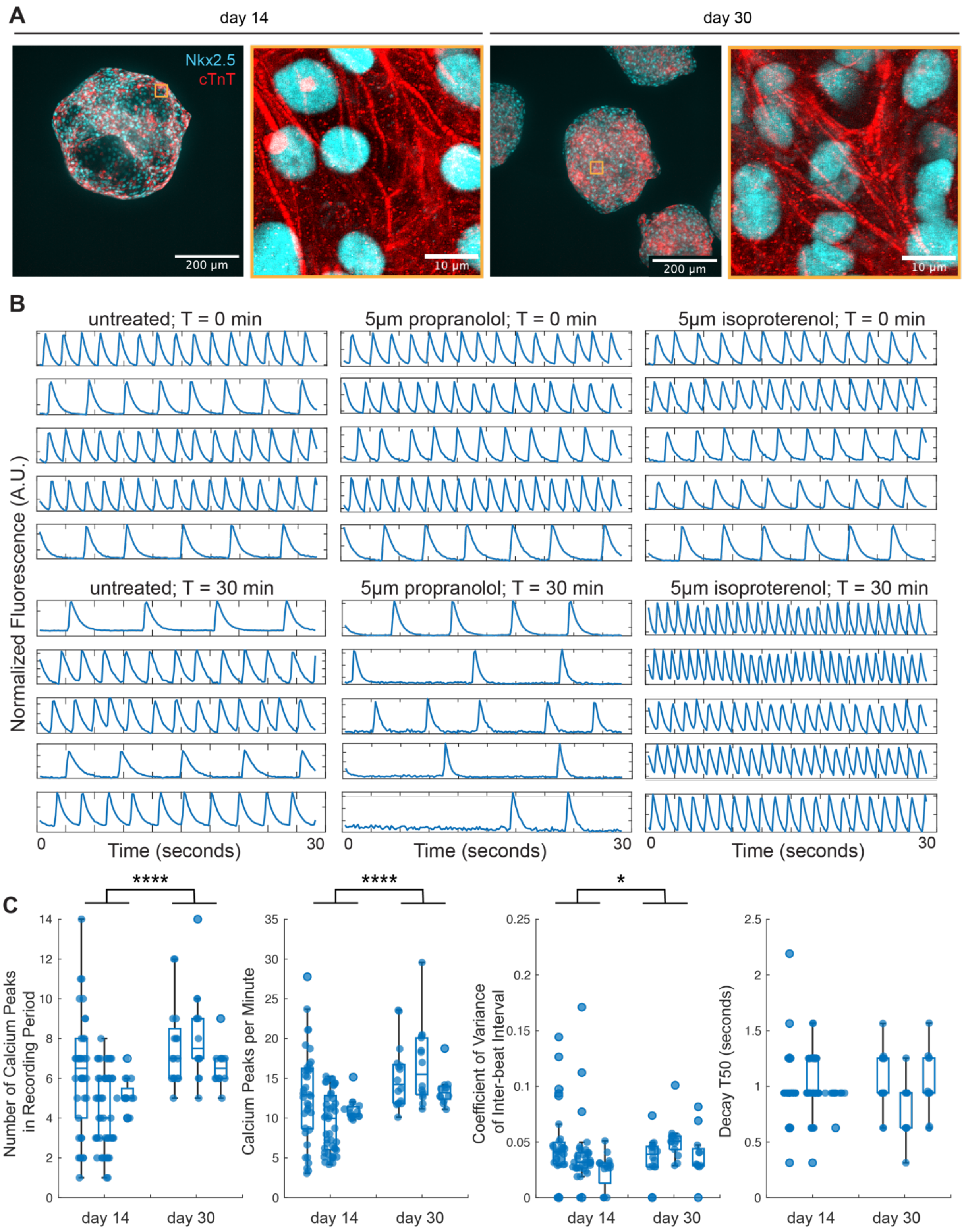


**Figure S9. saRNA-based delivery of jRCaMP1b to observe cardiac differentiation and drug response. Related to Figure 4 and Methods. A.** Representative immunofluorescent imaging of jRCaMP1b expressing cardiac spheroids fixed at day 14 and day 30 of differentiation. **B**. Representative jRCaMP1b traces for control, propranolol, and isoproterenol treated groups before (T = 0 min) and after 30 minutes (T = 30 min) of exposure. All traces are taken at 300 ms exposure time and are normalized between 0 and 1 to standardize temporal comparisons. Related to Fig. 4E. **C**. Quantified calcium transient parameters for traces represented in Fig. 4G prior to 10x interpolation for display purposes. At 3.33 fps temporal resolution, parameters appear quantized in 300 ms intervals, but statistical trends remain consistent with up-sampled traces (as reported in Fig. 4G) between day 14 and day 30.

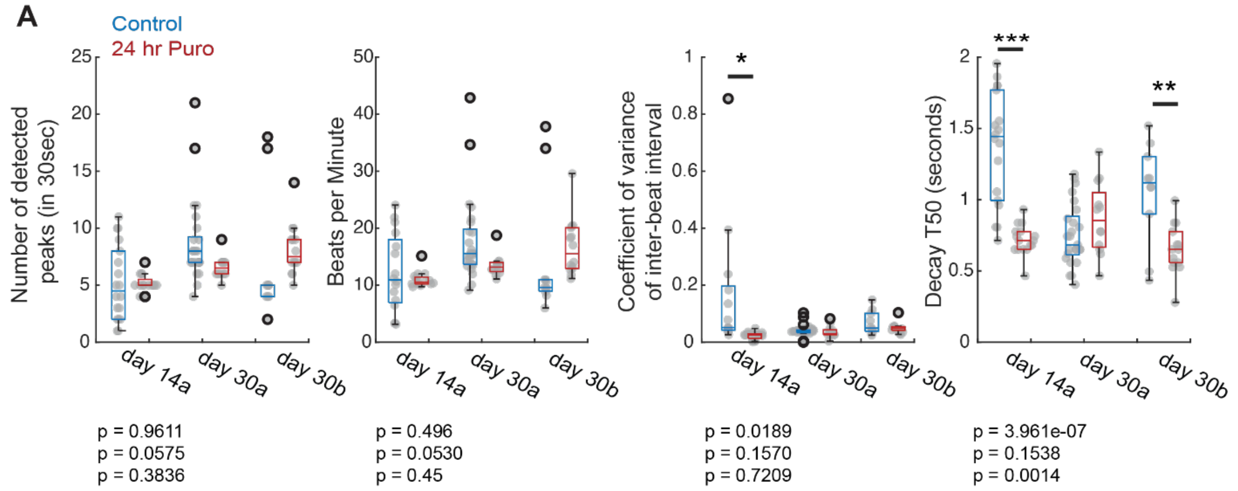


**Figure S10. Effect of 24-hour puromycin selection on cardiac spheroid differentiation. Related to Figure 4 and Methods. A.** Comparison of calculated calcium trace parameters between wells without initial selection and with 24-hour incubation with 1μg/mL puromycin following initial seeding into suspension. Day 14a and day 30a represent the same differentiation and day 30b is from an independent biological replicate. Statistical significance was calculated using one-way ANOVA analysis. From left to right in each plot N = 18, 16, 11, 14, 25, 12.

**Table S1. Sequences of oligonucleotides used for qPCR. Related to Methods.**

| Primer | Sequence (5'-3') |
|---|---|
| eGFP-Fwd | AGTCCGCCCTGAGCAAAGA |
| eGFP-Rev | TCCAGCAGGACCATGTGATC |
| mNGN2-Fwd | AGACGGTGCAGCGCATCAAGAA |
| mNGN2-Rev | AGCGTCTCGATCTTCGTGAGCT |
| BRN2-Fwd | ACCCGCTTTATCGAAGGCAA |
| BRN2-Rev | CCTCCATAACCTCCCCCAGA |
| ISL1-Fwd | AAGGTGGAGCTGCATTGGTTTG |
| ISL1-Rev | TAAACCAGCTACAGGACAGGCC |
| Nestin-Fwd | ACCCGCTTTATCGAAGGCAA |
| Nestin-Rev | AAGCTGAGGGAAGTCTTGGAGC |
| MAP2-Fwd | AGACTGCAGCTCTGCCTTTAG |
| MAP2-Rev | AGGCTGTAAGTAAATCTTCCTCC |
| TUBB3-Fwd | CAACCAGATCGGGGCCAAGTT |
| TUBB3-Rev | CCGAGTCGCCCACGTAGTT |
| GRIA4-Fwd | GGCCAGGGAATTGACATGGA |
| GRIA4-Rev | AACCAACCTTTCTAGGTCCTGTG |
| SYP-Fwd | ACCTCGGGACTCAACACCTCGG |
| SYP-Rev | GAACCACAGGTTGCCGACCCAG |
| VGLUT2-Fwd | GTAGACTGGCAACCACCTCC |
| VGLUT2-Rev | CCATTCCAAAGCTTCCGTAGAC |
| RPL37A-Fwd | GTGGTTCCTGCATGAAGACAGTG |
| RPL37A-Rev | TTCTGATGGCGGACTTTACCG |
